# Generalization reveals asymmetric and history-dependent control networks for multi-finger dexterous movements

**DOI:** 10.1101/2022.06.07.495015

**Authors:** Gili Kamara, Ohad Rajchert, Firas Mawase

**Affiliations:** Faculty of Biomedical Engineering, Technion – Israel Institute of Technology

**Author notes:** Correspondence to: Firas Mawase, Faculty of Biomedical Engineering Technion – Israel Institute of Technology.

**Keywords:** Finger dexterity, flexion, extension, generalization, finger individuation.

## Abstract

Finger dexterity, fundamental in our daily lives, is manifested by the generation of multi-finger and multi-directional patterns of muscles activity during various motor tasks, and further, by the generalization of learning in one context to other contexts. Tying shoelaces, for example, requires precise coordination of multiple fingers, some active primarily in the flexion direction, others in the extension direction, and some immobile. Nevertheless, whether the control processes of these actions are independent or interact and potentially generalize across each other, remains unclear. In a set of experiments, we sought to characterize the behavioral principles underlying the control process, learning and generalization of dexterous extension and flexion movements. We developed an isometric dexterity task that precisely measures dexterity in terms of finger individuation, force accuracy and temporal synchronization during finger flexion and extension. First, we investigated learning and generalization abilities across flexion and extension directions, both within and across hands. To do so, two groups of participants were trained for 3 days in either the flexion or extension direction. We found improvement in all dexterity measures in both groups following training, though finger extension generally exhibited inferior dexterity. Surprisingly, while the newly acquired finger extension abilities generalized to the untrained flexion direction, the newly acquired finger flexion abilities did not generalize to the untrained extension direction. Generalization biases of the finger flexion direction were also evident in the untrained hand. Next, we examined whether the asymmetric generalization pattern of multi-finger dexterous movements was history dependent. We thus recruited skilled musicians who showed increased baseline levels of dexterity in both directions and found that the degree to which learning generalizes between two contexts was affected by prior experience. Overall, our data indicate that control of multi-digit dexterous patterns is direction-specific in humans, supporting the hypothesis that control circuits for learning of finger flexion and extension are overlapped in that they partially, but asymmetrically, transfer between directions. This ability, however, is modular as it depends on hand use and the history of prior training.

## Introduction

Playing a piano, or simply tying shoelaces, requires precise coordination of multiple fingers, some active in flexion direction, others in the extension and/or other directions and some stay immobile. The ability of the sensorimotor system to control these and other dexterous movements is fundamental in human daily life and in the survival of higher mammals endowed with some degree of finger dexterity. This extraordinary dexterous behavior is particularly manifested by the way we generate and use specific patterns of muscles activity of multiple fingers when learning complex motor tasks, and further when we generalize what has been learned in one context to other contexts^1, 2^. Successful coordination of movements with different effectors is particularly evident when observing the intricate multi-finger combinations of movements used by skilled musicians while playing their instruments. Furthermore, when learning to type or to play musical instruments, such as the piano or violin, individuals initially experience great difficulty in simultaneously producing the appropriate movements across multiple effector muscles. These musicians must then learn to produce appropriate forces to flex some fingers while extending or maintaining others immobile with the proper orientation, with the optimal configuration varying across different types of keystrokes.

Simultaneous flexion and extension is required in most hand functions, and thus the success in performing these functions depends on the ability to precisely coactivate flexor and extensors finger muscles of both hands. The coexistence of control signals of flexion and extension movements raises the question whether the control processes of these actions interact and potentially generalize between each other. It is presumed that the extent of circuit sharing between different behaviors depends on the similarities in the patterns of muscle activation and the state of the limb, such as the effector(s) used or the resultant limb kinematics or dynamics^3^. In the case of multi-finger flexor versus extensor movements, the temporal sequence of muscle activations and the organization of muscle synergies differ markedly, yet the effector (i.e., fingers) and limb kinematics (e.g., velocity) are quite similar. Therefore, if the learning circuit encodes only direction-specific muscle patterns, then learning might not be shared and generalized across directions; but if it encodes effector or limb kinematics or dynamics, then learning might be shared. To date, the exact relationship between learning and generalization between flexors and extensors of multi-finger movements is still unclear.

One possibility is that learning to control finger flexors is independent from, and does not interact with, learning to control finger extensors (*independent control hypothesis,* **Figure 1D**). Under this hypothesis, learning to produce a particular multi-finger configuration in the flexion direction (e.g., simultaneously flexing the right ring finger and right middle finger) should not generalize to the extension direction with the same fingers (e.g., simultaneously extending the right ring finger and right middle finger), and vice versa. If learning is based on direction and muscle-specific information, the benefits of practice should not generalize to similar multi-finger patterns in the extension direction. In this case, the task could be thought of as requiring the learning of novel finger dynamics restricted to the direction within the practiced hand. Support for the independent control hypothesis includes a recent high-resolution functional imagining study in human participants that reported evidence of two spatially distinct finger maps for finger flexion versus finger extension in the primary motor cortex (M1)^4^.

**Figure 1.**
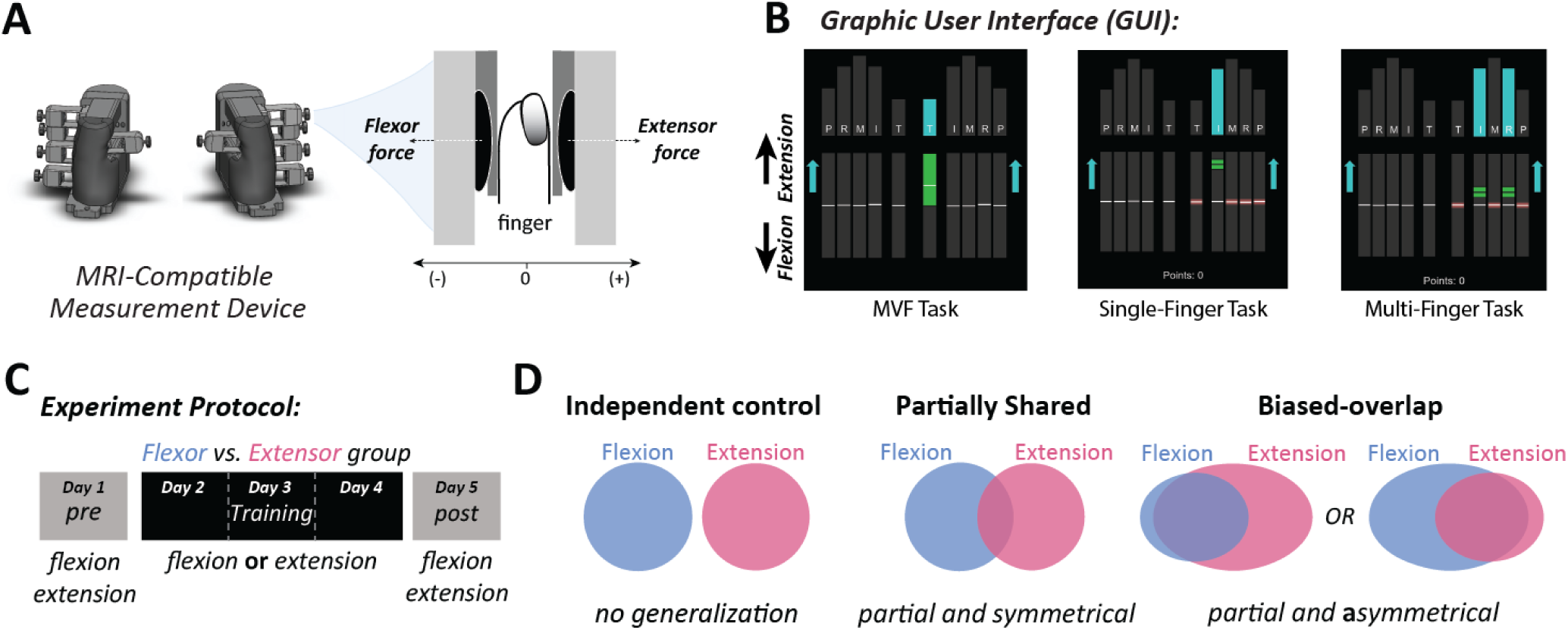
– Experimental setup, protocol, and research hypothesis. (A) The ergonomic device used to measure the isometric finger flexion and extension forces, using Honeywell force sensors as seen in the top view on the right (B) the Graphic User Interface (GUI) used in the experiment. The diagram on the left shows the GUI during the MVF task, specifically during the MVF measurement of right thumb extension. The middle diagram shows the GUI during an individuation (single-finger) task in which participants were asked to extend their right index finger to 75% of MVF. The diagram on the right shows the GUI during a chord (multi-finger) task in which participants were asked to extend their right index and ring finger simultaneously to 25% of MVF. (C) The experiment protocol was partitioned to three sections: pre-training testing, training, and post-training testing. All 4 movement types (right and left hands, both flexion and extension) were done on the testing days, though only right hand flexion or extension was done during the training, according to the group type. (D) Represents the distinct types of control patterns and associated prediction of each process as described in the hypothesis.

Another possibility is that learning in the flexion direction is partially shared with learning in the extension direction (*partially shared hypothesis*), and vice versa. One prediction of this interactive relationship hypothesis is that learning may not be restricted to the encoding of a specific direction, but rather might involve the encoding of the effector used (e.g., moving the ring and little fingers, regardless of the movement direction), and to some extent, the dynamic involved in the task (that is, applying %50 of maximum force), and that this generalizes across directions. In this view, multi-finger flexion-extension coordination would allow rapid and flexible alternation of finger movements in fine-motor control during dexterous movements. Support for the shared information between different muscle groups can be seen by the finding that primates’ single neurons in M1 receive common sensory input from the shoulder and elbow joints, but the output is largely specific to shoulder movements, potentially to allow rapid corrective responses to mechanical arm perturbation^5^. The shared hypothesis also makes an interesting prediction about the symmetricity of the generalization functions between flexion and extension movements. That is, the improvement in learning to the opposite direction is symmetric regardless of the trained direction.

While it may be intuitive to hypothesize symmetric generalization between flexion and extension, and vice versa, this is not necessarily true. First, pre-existing bias of flexion is evident by the ability to more precisely control movements of finger flexors compared to extensors^6^. Second, stroke survivors with cortical lesion who, despite regaining good flexion-based grasp, have very weak finger and wrist extensors that severely hamper hand opening^7–12^.

Third, a recent neurophysiological study showed increased representation of finger flexion, but weaker representation of finger extension, following micro-stimulation of the human motor cortex^13^. Thus, we proposed an alternative and third hypothesis (*biased-overlap hypothesis*) in which generalization across directions is asymmetrical. This hypothesis predicts different generalization patterns across directions and hands that depends on the trained direction.

We further examined the modularity of dexterity generalization and examined the effect of prior finger experience on learning and generalization of finger dexterity. We thus recruited a group of participants with extensive musical training that underwent a similar protocol to the other groups. Skilled pianists, for example, have higher levels of finger individuation than non-musicians^14, 15^, which is probably acquired through extensive training^16–20^. Although individuation and keystroke timing accuracy in skilled pianists are relatively invariant across a wide range of tempi^21^, more variability in timing occurs when less individuated digits play notes^22^. When playing, skilled pianists use lower impulse and force to achieve the same volume across keys (indicating higher energy efficiency) compared to novices^23^. Musical performance thus combines high levels of finger dexterity with enhanced individuation, precise control of force and temporal synchronization, not only in producing single notes, but also in producing precisely controlled multi-finger chords in flexion-extension directions. Whether this extensive experience in controlling fine-motor finger movements can affect across-direction generalization patterns remains unknown.

We therefore sought to characterize the behavioral principles underlying the control process, learning ability and generalization of single and multi-finger dexterous movements in the flexion and extension directions, via a novel finger dexterity task. We decomposed dexterity into its underlying components: *finger individuation,* defined as the ability to independently move the instructed finger(s), either in a flexion or extension direction, while keeping the uninstructed fingers immobile (minimizing enslaving), *force control,* defined as the ability to generate accurate force of the instructed finger(s), and *temporal synchronization,* defined as the ability to produce simultaneous movement between instructed finger(s).

We found evidence in support of a biased-overlap control network for finger flexion vs. extension. Following multiday training of a multi-finger dexterity task, we trained separate groups for 3 days in either the flexion or extension direction. We observed improvement in all dexterity measures in both groups, although inferior dexterity in the extension direction. Interestingly, we observed asymmetric generalization across directions. That is, learning generalized from the extension to flexion direction but not from flexion to extension of the trained hand, and that the generalization was only to the flexion direction of the untrained hand. The group of musicians, that was recruited to explore the effect of prior history of finger experience on generalization, exhibited much stronger and directionally unbiased generalization to the untrained direction of either hand. Together, these findings corroborate with our hypothesis that controlling multi-finger dexterous patterns is direction-specific and history-dependent in humans, and that learning is dissimilarly transferred between directions, supporting the hypothesis that neural control circuits for learning of finger flexion and extension interact and partially, but asymmetrically, overlap in the sensorimotor system.

## Methods

### Participants

In total, 52 right-handed participants (31 female), aged 25.7±3.7 years (mean ± STD), were recruited and given monetary compensation for their participation (300₪ ≈ $90). Approval of the local Ethics Committee was obtained, and informed consent was given by all participants prior to their inclusion. Participants’ handedness was evaluated using the Edinburgh Handedness Inventory 10 item version^24^. All participants were deemed fully capable in terms of motor abilities, with no history of brain damage or motor impairments affecting finger movements. Naïve participants with no musical history were divided into 3 separate groups based on the trained direction (flexion vs. extension): Flexor group (n=13) who trained in finger flexion, Extensor group (n=13) who trained in finger extension, and a control group (n=8) which performed two testing sessions 4 days apart, without undergoing training in between the tests. In addition, to investigate the effect of the history of prior finger dexterity skills on learning and generalization of finger dexterity, we recruited musicians (n=13, Musician group) that trained in the extension direction and examined the generalization pattern (post-training) on both directions. Non-musicians were considered as those with no more than 6 consecutive months of formal and/or informal musical training, whereas musicians had at least 4 years of formal musical training. Five participants were excluded from analysis due to incomplete or missing data, or dropout.

### User interface and data acquisition

The study utilized an adjustable ergonomic device that measured isometric finger forces, with a force sensor (FSG-020WNPB, Honeywell^®^; dynamic range 0-20 N) below each fingertip to measure flexion forces and a sensor above each fingertip to measure extension forces (see **Figure 1A**). During the study, participants placed both hands in a neutral posture inside the device. Analog force signals were digitized and sampled at 250 Hz (using NI USB-6211 data acquisition), and then integrated with a customized MATLAB script (The MathWorks, Inc., Natick, MA, ver. R2020b) enabling live measurements, presentation, and analysis. Using Psychtoolbox^25^, visual stimuli presented on a computer screen signaled to the participants which fingers they were required to move and at what force. The target forces were normalized to 25, 50, and 75% of each participant’s maximum voluntary force (MVF) for each finger.

During testing and training, participants were instructed to move only the relevant fingers until reaching the appropriate target forces while maintaining the uninstructed fingers at rest. A force tolerance around each target force represented the target zone in green, whereas uninstructed fingers had a red zone of ±5% MVF around 0 N. For example, **Figure 1B** shows a trial in which participants were instructed to extend their thumb and index fingers to 25% of MVF while maintaining the uninstructed fingers immobile. Participants were incentivized with a point and a success sound when they completed a trial correctly (i.e., all instructed fingers were within the green zones and all uninstructed fingers were within the red zone). Points were aggregated during each block and the total point count from all blocks was displayed at the end of each block. Participants were explicitly told that this feedback was not related to the compensation for their participation in the experiment.

### Study design

The primary aim of this study was to characterize the learning and generalization of finger dexterity in the flexion and extension directions. Therefore, two different cohorts of participants (Flexor and Extensor) were trained with the dominant right hand over three sessions (days 2– 4), in the flexion or extension direction, respectively. We first quantified the learning effect in multiple dexterity measures within the trained direction. We then quantified the generalization of learning effects to the untrained direction (directional generalization) and to the untrained hand (lateral generalization) during the post-training tests conducted after training (on day 5). An additional group of non-musical participants was used as a control group which did not partake in training but only performed the tests on day 1 and day 5. A secondary aim was to investigate whether learning and generalization are affected by the history of prior finger skill practiced over years. To do so, we recruited a cohort of professional musicians with years of experience playing musical instruments (e.g., piano, violin) that undergo similar protocol.

More specifically, the study spanned 5 consecutive days (see **Figure 1C**). On the first day (pre-training), the participants’ MVF and baseline measurements were acquired. On days 2, 3, and 4 the participants performed the training according to their group type. On the fifth day (post-training), the participants underwent the same measurements as on the first day. ***Day 1 (pre-training)*** – Following an explanation of the visual stimuli and a short demonstration of the hand sensors, the hand device was adjusted according to the participants’ hand dimensions. The MVF of each finger was obtained using the ‘Finger strength task’ (Figure 1B). Next, initial baseline measurements were obtained using the ‘Individuation and chord testing task’. ***Day 2, 3, 4*** – Participants performed the ‘Training task’, according to their group type. ***Day 5 (post-training)*** – Participants performed the ‘Individuation and chord testing task’, as in day 1, and received compensation for their participation.

#### Trial and block design

The study was built in a session, block and trial design. Each session was separated into multiple blocks, and each block contained multiple trials. Participants were allowed to remove their hands from the devices between blocks. Each trial in the testing and training tasks consisted of the same procedure: The GUI signaled which finger is to move 750 ms prior to displaying the green target zones, and participants had 5 seconds to successfully enter the green zones. If reached, participants were instructed to hold the force and stay within the green zone for two seconds. If all instructed fingers were within the green zones and all the uninstructed fingers were within the red zones, then the trial was considered successful, and the participants were awarded with a point and a ‘success’ tone. The next trial began automatically after 250 ms (inter-trial interval, ITI).

### Dexterity testing and training tasks

All tasks were performed when the participant was seated facing a computer screen with both hands inside the hand devices. Participants could remove their hands from the device between blocks. Prior to the beginning of each task, participants were provided with an overview of the task and its requirements. The tasks used are similar to those used in previous works^26, 27^. In order to accurately measure the isometric forces during activity while considering forces produced by the hand placement and finger weights, we subtracted forces measured by the sensors while participants were in a neutral position and at rest. This zeroing procedure was performed before each block.

### Finger strength task

This was the first task performed by the participants in order to find the MVF of each finger in each direction. During each trial, participants were asked to produce as much force as they could in a certain finger and direction and to maintain that force for three seconds (**Figure 1B**, left panel). Here, participants were not limited to using only the instructed finger to produce the maximal force. Due to limitations in the force capability of the sensors, participants were allowed to produce forces up to 14 N. If the force produced exceeded 14 N, then the green target zone turned red, and participants were asked to reduce their force slightly. Forces larger than 14 N mainly occurred during thumb and index flexion, if at all, so this limitation did not affect all participants nor all MVF values.

The order in which the MVF values were measured was right hand (RH) flexion, RH extension, left hand (LH) flexion, and LH extension, in order from thumb to little finger. The maximum force of each trial was calculated as the 95^th^ percentile of the finger’s force data for the trial. Each movement was performed twice and the maximum between the two repetitions was selected as the relevant MVF value.

### Single finger individuation and multi-finger chord tasks

This task was performed on the first and last day of the study, to provide the baseline (pre-training) and final (post-training) learning abilities, respectively. Participants were instructed to move only the instructed finger(s) while maintaining the other fingers at rest. This task consisted of 388 trials in 8 separate blocks, each of which tested a different subgroup of movement: RH Individuation - Flexion, RH Individuation - Extension, LH Individuation - Flexion, LH Individuation - Extension, RH Chords - Flexion, RH Chords - Extension, LH Chords - Flexion, and LH Chords - Extension.

The individuation blocks consisted of 3 repetitions of three force levels (25%, 50%, and 75% of MVF) for each finger, totaling at 45 trials per block. Trials were performed in order from thumb to little finger. The chord blocks consisted of two repetitions of the 25% force level in the remaining 26 finger combinations (i.e., a total of 31 possible combinations in one hand, including 5 single finger movements and 26 multiple-finger combinations), totaling at 52 trials per block. Chord trials were performed in an intuitive order, starting from two-finger combinations and ending at all five fingers. The force tolerance around each target force was set at 10% of MVF. This task took roughly 60-75 minutes to complete. Participants of all groups performed identical pre- and post-training tests.

### Training task

The training task was performed on days 2,3 and 4 of the study and included a total of 310 trials in 5 blocks. Each block contained two repetitions of all 31 finger combinations at the 25% force level, presented in random order. Training was done only on the right hand. The direction of the training depended on the group to which the participant belonged: ‘Flexors’ (non-musicians, flexion); ‘Extensors’ (non-musicians, extension); and ‘Musicians’ (extension). The force tolerance around the target forces decreased by 85% per day (8.5%, 7.2%, and 6.1% of MVF, respectively). This task took approximately 50-60 minutes to complete.

### Measurable metrics

Prior to extracting metrics from the raw data, various preprocessing analyses were implemented. (1) *Low-pass filtering* - The force data of each sensor was filtered using a gaussian filter (over a 16 ms time window) to eliminate the electrical noise from the sensors. (2) *Baseline-correction* - The forces present at the beginning of each trial (baseline forces) were calculated over the initial 1.5 seconds of the trial in three segments (0-0.5, 0.5-1.0, 1.0-1.5 seconds). In order to verify that the finger was at rest, the standard deviation of each segment was calculated. If the standard deviation of the segment with the lowest standard deviation was less than 0.2, then the mean force of that segment was chosen as the baseline force. The baseline forces, if available, were then subtracted from the data, resulting in modified forces which start at 0N for each trial. Sample processed trial data of single finger movement can be seen in **Figure 2B**, and sample processed trial data of multi-finger movement can be seen in **Figure 2C**. Following the preprocessing stage, the peak force of each instructed finger was found and was used to find the movement onset and end times. The onset and offset timestamps were classified as the first and last instances in which the force passed 50% of the peak force, respectively. If there was more than one instructed finger, the longest range was selected (see onset times in Figure 2F).

**Figure 2.**
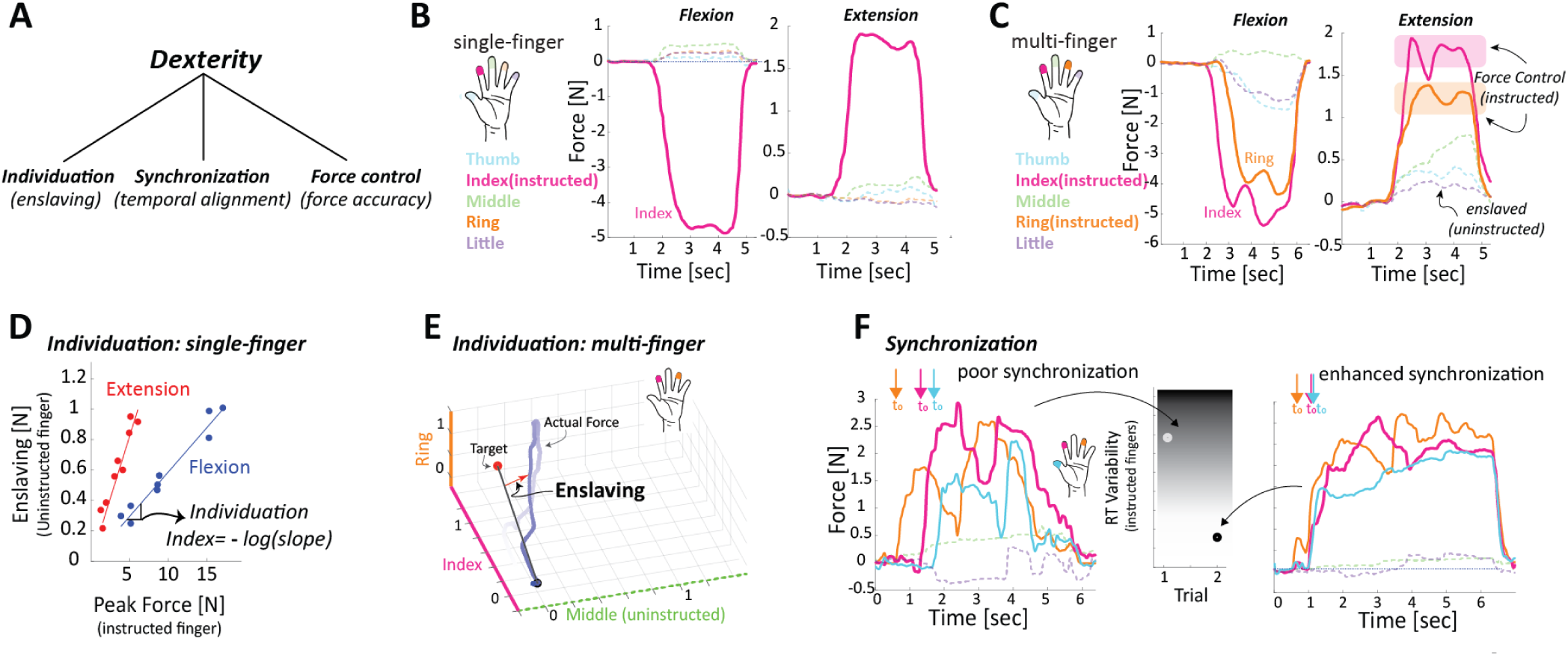
– Visual descriptions of measurable metrics. (A) Dexterity is characterized by 3 main quantifiable components. (B) Example of a single-finger flexion and extension trial, specifically isolated finger movements. Flexion forces were assigned negative values and extension forces had positive value. Force targets were set according to each finger’s MVF. (C) Examples of a multi-finger flexion and extension trial, specifically index and ring finger movements. Accuracy, or force control, was calculated for the instructed fingers’ force profiles around the target forces, as seen in the highlighted regions. Enslaving was calculated over the same timeframe from the uninstructed fingers’ forces (dashed lines). (D) The Individuation Index of each finger was calculated as the -log of the slope between the peak active force and the uninstructed finger deviation. This graph contains the 9 extension (red) and 9 flexion (blue) trials (3 repetitions of 3 force levels) of a single finger. (E) A 3D representation of the enslaving which occurs during multi-finger movements. (F) Poor temporal synchronization can be seen in the left trial of multi-finger trials, compared to a trial with enhanced temporal synchronization. The movement onset of the different instructed fingers is noted with arrows.

### Individuation Index (II)

One of the main components of finger dexterity is the ability to individuate the fingers. In order to quantify this ability and test how it is altered and potentially generalized following training, a metric called Individuation Index (*II*) was calculated and compared over the duration of the study^27^. The *II* describes the relationship between the forces generated by the instructed finger and the uninstructed fingers. The *II* was derived from the mean deviations of the uninstructed fingers and the peak force of the instructed finger in each specific trial. The mean deviation of the uninstructed fingers (*meanDevU*) was calculated using the following equation:

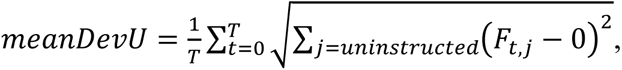

where the index j denotes the j^th^ uninstructed finger, T and t represent time, Ft,j is the j^th^ finger’s force level at time t, and the deviation was calculated from F=0N. The timespan evaluated for this calculation was between the movement onset and end times (see arrow in **Figure 2C**). The peak force and mean deviation values of the various trials for the same finger present a positive linear relationship, with increasing uninstructed finger deviation for increased peak instructed finger force. The *II* of each finger was calculated as the negative log of the slope, and the *II* of each hand was the average of the finger *II* values (see **Figure 2D**). The higher the *II* value, the better the individuation ability.

### Force control of the instructed fingers

Dexterity can also be described in terms of force accuracy of the instructed fingers. The ability to generate accurate force of the instructed finger(s) was quantified by measures of deviation from the target (i.e., *meanDevI*), using the following equation:

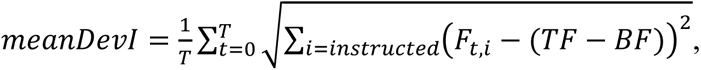

where the index i denotes the i^th^ uninstructed finger, T and t represent time, Ft,i is the i^th^ finger’s force level at time t, TF is the predefined target force, and BF is the baseline force calculated during the preprocessing stage. The timeframe for the calculation was the movement onset and offset times as defined previously (see pink and orange regions in **Figure 2C**)

### Temporal synchronization

In the time domain, one of the main principles underlying the control process of multi-finger dexterous movements is how participants synchronize their force patterns when presented with targets incorporating multiple fingers. Synchronization is defined as the temporal alignment between active fingers and is calculated as variance (e.g., standard deviation) of the response, or reaction time (RT), of the instructed fingers. Low RT variability indicates enhanced synchronization between the instructed fingers (right panel, **Figure 2F**), whereas high variability indicates reduced synchronization (left panel, **Figure 2F**). RT of each finger is defined as the time interval between the “Go” cue and the force initiation (detected as the time at which the force signal crossed the threshold of 10% of peak force). In chord targets, the overall RT was considered the reaction of the first finger to surpass the 10% threshold.

### Statistical analysis

The statistical analysis was performed using Matlab software (MathWorks) and Prism software (GraphPad). To determine the effect of learning in force control (i.e., deviation of the instructed fingers), enslaving (i.e., deviation of the uninstructed fingers) and synchronization, we performed for each group two-tailed paired t-tests between the first block of the training (i.e., day 2) and the last block of the training (i.e., day 4). To determine the across-direction generalization, we then used separate (for each group) 2-way repeated-measures ANOVA (2-way RM-ANOVA) to assess differences in individuation index, force control, and synchronization of the trained hand, with factors of time (pre vs. post training) and direction (flexion vs. extension). For across-hand generalization statistics, we also used separate 2-way RM-ANOVA with factors of time (pre vs. post training) and direction (flexion vs. extension). When significant differences were identified, post hoc analysis was conducted using the Holm-Šídák paired t-test for correcting multiple comparisons. In all comparisons, the significance level was set at 0.05.

## Results

### Within direction learning of flexion and extension multi-finger actions

During the training phase of the study (days 2, 3, and 4), participants performed each of the 31 finger combinations a total of 10 times during 5 blocks (2 repetitions of each combination in each block) in which the combinations were presented in a pseudo-random order. Each group trained only in its respective direction, and the following values represent within-direction learning metrics during the training period.

### Accuracy as a measure of force control

The accuracy of each trial was described by the *meanDevI*. The lower the *meanDevI*, the more accurate the instructed finger(s) were in achieving the target forces. The improvement in accuracy during the training period can be seen by the decreasing trend in **Figure 3A**. Because flexion motions generally had a larger maximal force than extension motions, the deviation values were normalized by the fingers’ MVF in order to enable comparison between the two groups and their respective directions. The daily averages of % MVF (±SE) of the flexor group were 8.73 (±0.14)%, 7.75 (±0.07)%, and 6.90 (±0.06)% for days 2-4, respectively, with an improvement of 2.68% (i.e., 9.50%-6.82%) between the first and last blocks; the daily averages of % MVF of the extensor group were 11.27 (±0.17)%, 9.94 (±0.13)%, and 9.40 (±0.10)% for days 2-4, respectively, with an improvement of 2.70% (i.e., 12.16%-9.46%) between the first and last blocks. The flexor group was consistently more accurate than the extensor group, though both groups had similar trends of improvement. This can be seen in the 2-Way (time vs. group) RM-ANOVA, which showed a significant time effect as a result of the training (F(1,23)=87.28, p<0.0001), as well as a significant difference between the two groups (F(1,23)=22.65, p<0.0001), though there was no significant interaction between the time and group factors (F(1,23)=2.865, p=0.104).

**Figure 3.**
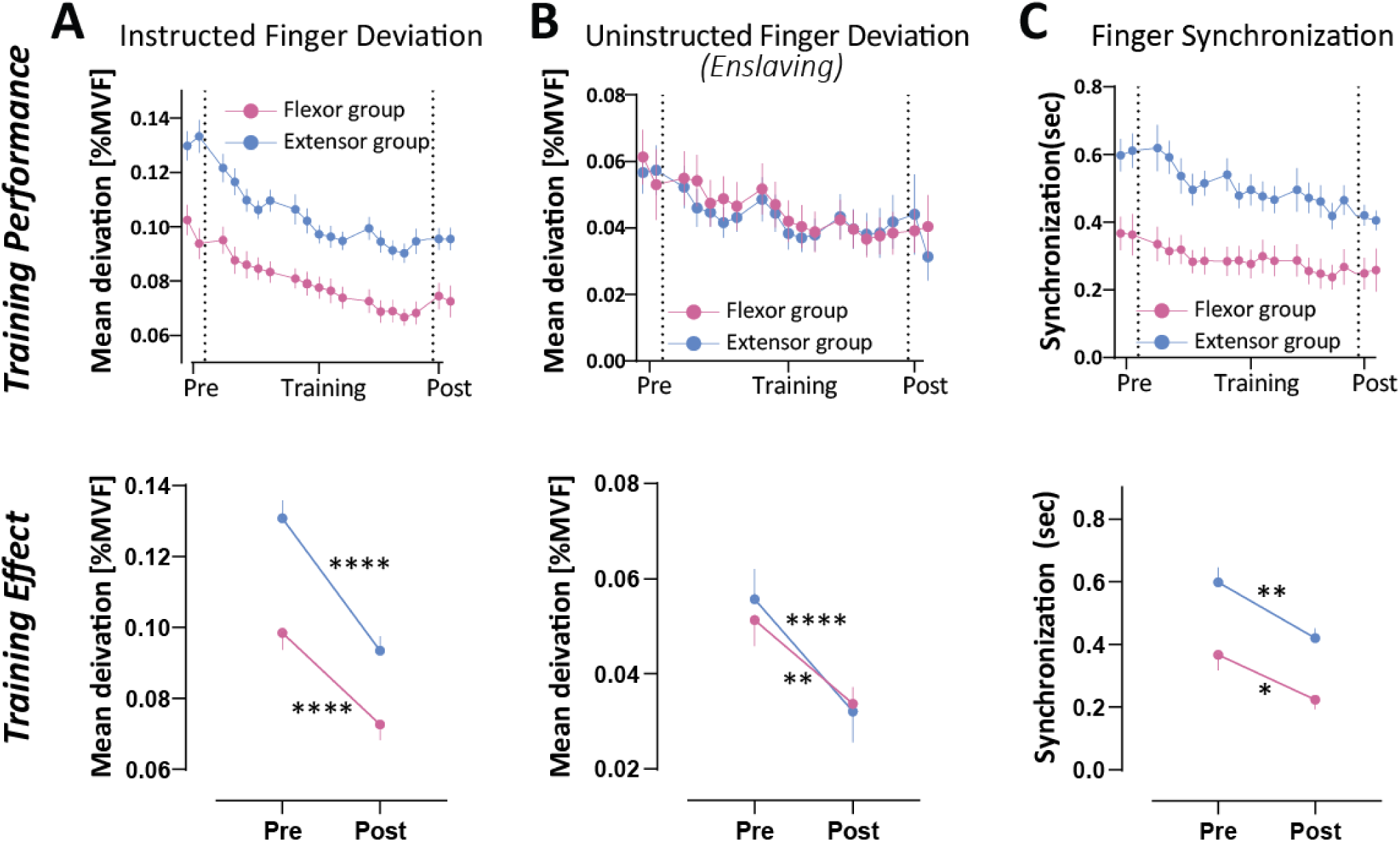
– Performance during the training period and training effect in the various metrics on the trained hand and trained direction of both flexor and extensor groups. Training Performance metrics represent the average performance in each of the 19 blocks; Pre and Post values are the average performance during the averaged values of the two testing blocks. (A) The force control metric is depicted using the ‘Instructed Finger Deviation’, or MeanDevI, normalized by the MVF. (B) The Uninstructed Finger Deviation, or MeanDevU, normalized by the MVF. (C) The temporal synchronization between fingers.

### Uninstructed finger deviation (enslaving)

The uninstructed finger deviation of each trial was described by the *meanDevU*. The lower the *meanDevU*, the less there was enslaving or movement in the uninstructed fingers. The improvement in abilities can be seen by the trend of decreasing deviation during the training period (**Figure 3B**). Here again, values were normalized by the fingers’ MVF in order to enable comparison between flexion and extension movements. The daily averages of % MVF (±SE) of the flexor group were 4.43 (±0.11)%, 3.83 (±0.15)%, and 3.39 (±0.06)% for days 2-4, respectively, with an improvement of 1.52% (i.e., 4.82%-3.31%) between the first and last blocks; the daily averages of the extensor group were 4.56 (±0.11)%, 4.13 (±0.14)%, and 4.03 (±0.06)% for days 2-4, respectively, with an improvement of 1.05% (i.e., 5.23%-4.18%) between the first and last blocks. Two-way (time vs. group) RM-ANOVA showed a significant time effect as a result of the training (F(1,22)=40.28, p<0.0001), though there was no significant difference between the two groups (F(1,23)=0.1639, p=0.8993) nor any significant interaction between the time and group factors (F(1,22)=1.073, p=0.3115). Though the flexor and extensor groups had similar initial values, the flexor group had a larger improvement during the training.

### Reaction time

During multi-finger targets, the reaction time was selected as the minimal timestamp of the various instructed fingers. Due to the randomized order of finger combinations, participants could not anticipate and/or plan their motion prior to the visual cues. The daily averages of RT (±SE) of the flexor group were 707.9 (±18.3) ms, 590.4 (±11.6) ms, and 555.5 (±9.0) ms for days 2-4, respectively, with an improvement of 245.1ms (i.e., 791.9-546.8ms) between the first and last blocks; the daily RT averages of the extensor group were 790.4 (±22.8) ms, 692.9 (±12.2) ms, and 643.7 (±10.8) ms for days 2-4, respectively, with an improvement of 282.4 ms (i.e, 924.9-642.5) between the first and last blocks. Two-way (time vs group) RM-ANOVA showed a significant time effect as a result of the training (F(1,22)=27.09, p<0.0001), though there was no significant difference between the two groups (F(1,22)=0.618, p=0.4401) nor any significant interaction between the time and group factors (F(1,22)=0.861, p=0.3635), suggesting that both groups improved their RT in a relatively comparable manner.

### Temporal synchronization during multi-finger actions

The degree of synchronization among multi-finger actions was described by the standard deviation between the various instructed fingers’ reaction times. The lower the standard deviation, the more synchronous the motion. The daily RT averages (±SE) of the flexor group were 355.5 (±5.0) ms, 306.0 (±5.7) ms, and 286.2 (±0.3) ms for days 2-4, respectively, with an improvement of 108.7 ms (i.e., 367.4-258.7 ms) between the first and last blocks; the daily RT averages of the extensor group were 609.9 (±3.0) ms, 542.0 (±13.4) ms, and 511.9 (±8.7) ms for days 2-4, respectively, with an improvement of 192.6 ms (i.e., 598.3-405.7) between the first and last blocks. In this metric, the flexor movement had significantly better temporal synchronization than the extension movement, though both groups showed improvement during the training. Two-way (time vs group) RM-ANOVA showed a significant time effect as a result of the training (F(1,22)=23.23, p<0.0001), as well as a significant difference between the two groups (F(1,22)=20.78, p=0.0002), though there was no significant interaction between the time and group factors (F(1,22)=0.2624, p=0.6136). **Figure 3C** shows the improvement in synchronization over the course of the training period for both groups.

### Individuation index

The individuation indices were measured on the first and last day of the study. The higher the II, the better the individuation ability (see Table 1 for full results). On the first day, the RH flexion overall hand II (±SE) average of the flexor group was 2.36 (±0.16). Following the training, the RH flexion overall hand II (±SE) average of the flexor group was 2.65 (±0.17), resulting in an improvement of Δ=0.29 (Figure 4A, flexion direction). Post-hoc two-tailed, paired sample t-test comparison between performance on Day 5 and Day 1 showed that training led to significant improvement in RH flexion II values (t(12)=3.717, p=0.003). Similarly, the extensor group also showed positive change in overall II following training (Figure 4B, extension direction). The RH extension overall hand II (±SE) pre-training was 2.25 (±0.17). Following the training, the RH extension overall hand II (±SE) was 2.58 (±0.14), resulting in an improvement of Δ=0.33 following the training. Post-hoc paired two-tailed, sample t-test comparison between performance on Day 5 and Day 1 showed that training led to significant improvement in RH extension II values (t(12)=3.006, p=0.011). These results show that both groups improved in their trained direction to a similar degree in terms of overall II.

**Figure 4.**
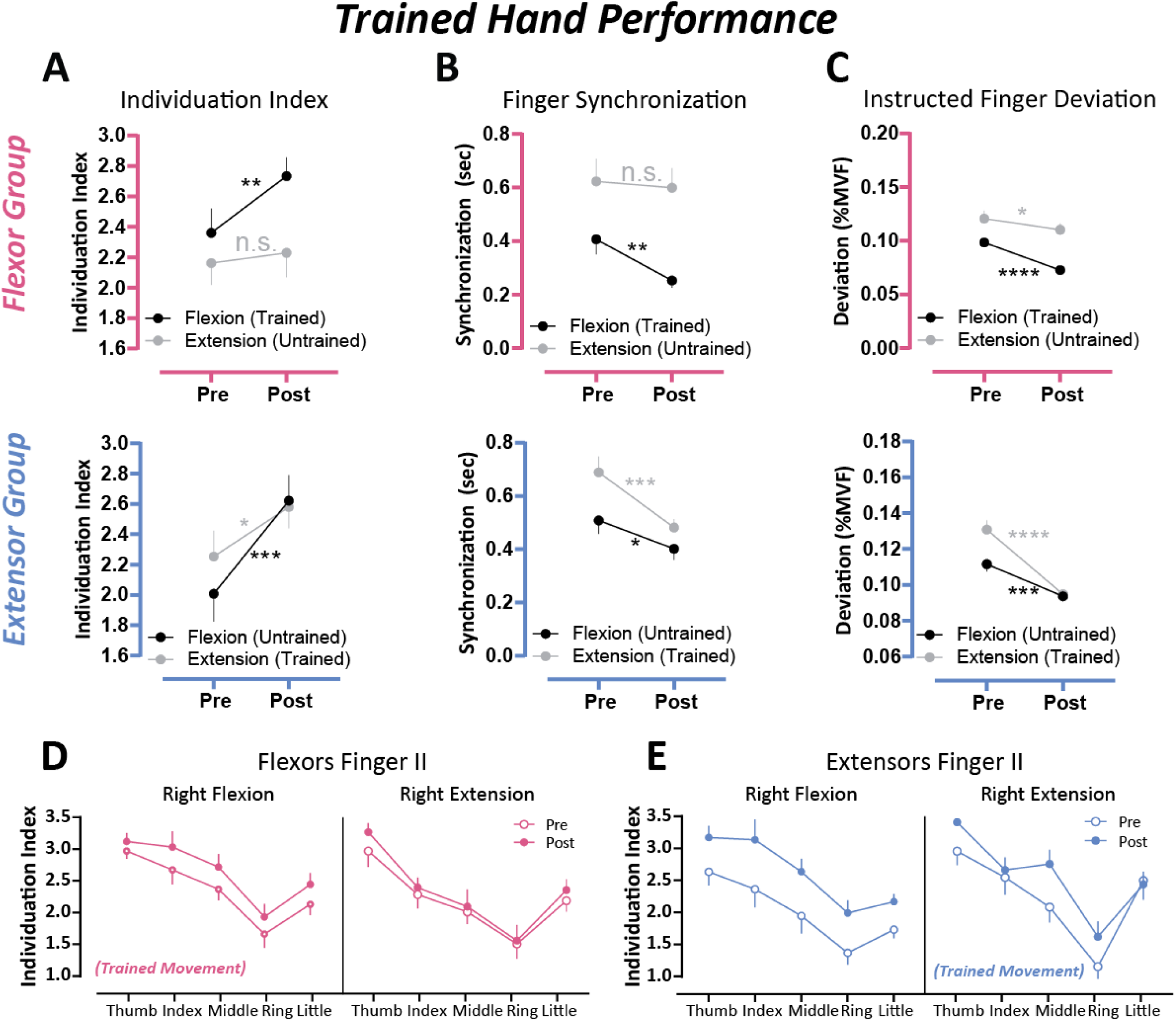
– Asymmetric generalization of finger dexterity measures across directions of the trained hand of both Flexor and Extensor groups. (A) Training effect on the values of Individuation Index. (B) Same as (A) but for changes in temporal synchronization between fingers. (C) Same as (A) but for force control, as calculated by MeanDevI, normalized by the MVF. (D) The Flexor group’s finger II values on RH flexion and RH extension, pre and post training. (E) The Extensor group’s finger II values on RH flexion and RH extension, pre and post training.

### Directional generalization

Movements in the opposite direction of the training were performed and measured only on the first and last day. The RH extension II (±SE) of the flexor group was 2.16 (±0.14) prior to the training, and 2.31 (±0.13) following the training period, resulting in an improvement of Δ=0.15 (Figure 4A, top panel). Two-way (time vs. direction) RM-ANOVA was performed for the flexor group, which showed a significant effect for both time (F(1,12)=6.40, p=0.0264) and direction parameters (F(1,12)=7.39, p=0.0187), and significant, yet marginal, interaction between the two (F(1,12)=4.43, p=0.0569). While the flexor group improved II in the trained flexion direction of the right hand, post-hoc two-tailed, paired sample t-test comparison between performance on Day 5 and Day 1 showed that training did not lead to significant improvement in RH extension II values (t(12)=1.085, p=0.299). The asymmetry between the flexion and extension IIs on the trained hand can be seen in detail in Figure 4D, which shows a rather consistent improvement among all fingers in the flexion motion, though only a slight difference in individual fingers’ II values for extension motion. For the extensor group, the RH flexion II (±SE) of the hand was 2.01 (±0.18) pre-training, and 2.62 (±0.17) post-training with an improvement of Δ=0.61 (Figure 4A, bottom panel). Two-way (time vs. direction) RM-ANOVA was performed for the extensor group, which showed a significant effect for the time parameter (F(1,12)=23.97, p=0.0004), though not for the direction parameter (F(1,12)=0.3926, p=0.5427), but marginal significant interaction between the two (F(1,12)=4.10, p=0.0657). Interestingly, post-hoc two-tailed, paired sample t-test comparison between performance on Day 5 and Day 1 showed that training led to significant improvement in RH flexion II values in the extensor group (t(12)=6.127, p<0.0001). The asymmetry between the flexion and extension IIs on the trained hand can be seen in detail in Figure 4E, which shows a rather consistent improvement among all fingers in the flexion motion, though inconsistent differences in extension motions. A comparison between Figure 4D-E further shows the effect of the trained direction on the learning and generalization abilities. These results depict an asymmetry in directional generalization, which will be elaborated on further in the discussion.

The average temporal synchronization of RH extension in the Flexor group prior to the training was 618.7 (±79.2) ms, and following training was 568.8 (±68.4) ms, resulting in an improvement of Δ=49.9 ms (Figure 4B, top panel). Two-way (time vs. direction) RM-ANOVA was performed for the flexor group, which showed a significant effect for the direction parameter (F(1,11)=19.10, p=0.0011), and a possible statistical trend for the time parameter (F(1,11)=3.793, p=0.0774). There was an interaction between the time and direction parameters (F(1,11)=7.043, p=0.0224). For the extensor group, the average RH flexion synchronization (±SE) prior to training was 519.0 (±52.9) ms, and following training was 412.3 (±42.1) ms, resulting in an improvement of Δ=106.7 ms (Figure 4B, bottom panel). Two-way (time vs. direction) RM-ANOVA was performed also for the extensor group, which showed a significant effect for both the time parameter (F(1,11)=20.44, p=0.0009) and the direction parameter (F(1,11)=14.15, p=0.0031), though there was no interaction between the two (F(1,11)=3.315, p=0.0960).

The average accuracy, measured as *meanDevI* normalized by MVF, of RH extension of the flexor group prior to the training was 0.120 (±0.0068), and following training was 0.110 (±0.006), resulting in an improvement of Δ=0.01 (Figure 4C, top panel). Two-way (time vs. direction) RM-ANOVA was performed for the flexor group, which showed a significant effect for both the time parameter (F(1,11)=16.97, p=0.0017) and the direction parameter (F(1,11)=32.23, p=0.0001), as well as an interaction between the two (F(1,11)=9.840, p=0.0095). For the extensor group, the average RH flexion accuracy (±SE) prior to training was 0.1123 (±0.0051), and decreased to 0.1109 (±0.0346) post-training, resulting in an improvement of Δ=0.0054 (Figure 4C, bottom panel). Two-way (time vs. direction) RM-ANOVA was also performed for the extensor group, which showed a significant effect for both the time parameter (F(1,11)=62.32, p<0.0001) and the direction parameter (F(1,11)=10.62, p=0.0068), as well as an interaction between the two (F(1,11)=11.49, p=0.0069).

### Across hands generalization

Movements of the opposite hand (left hand) were only performed on the first and last day as part of the testing tasks. Prior to the training, the average LH flexion II (±SE) was 2.13 (±0.15), and following training was 2.45 (±0.16), resulting in an improvement of Δ=0.32 (Figure 5A, top panel). Two-way (time vs. direction) RM-ANOVA was performed for the extensor group, which showed a significant effect for the time parameter (F(1,12)=7.30, p=0.0192), though not for the direction parameter (F(1,12)=0.61, p=0.4474), nor an interaction between the two (F(1,12)=3.09, p=0.1042). Post-hoc two-tailed, paired sample t-test comparison between performance on Day 5 and Day 1 showed that training led to significant improvement in left hand flexion II values (t(12)=4.103, p=0.003). For the extensor group, the average LH extension II (±SE) prior to training was 1.97 (±0.14), and increased to 2.10 (±0.14) post-training, resulting in an improvement of Δ=0.13 (Figure 5A, bottom panel). Two-way (time vs. direction) RM-ANOVA was also performed for the extensor group, which showed a significant effect for both time (F(1,12)=4.85, p=0.0478) and direction parameters (F(1,12)=6.85, p=0.0224), though no interaction between the two (F(1,12)=1.45, p=0.2503). Post-hoc two-tailed, paired sample t-test comparison between performance on Day 5 and Day 1 showed that training did not lead to significant improvement in LH extension II values (t(12)=1.556, p=0.146). Here again, asymmetry between the generalization patterns of the learned flexion and extension, due to the training, can be observed.

**Figure 5.**
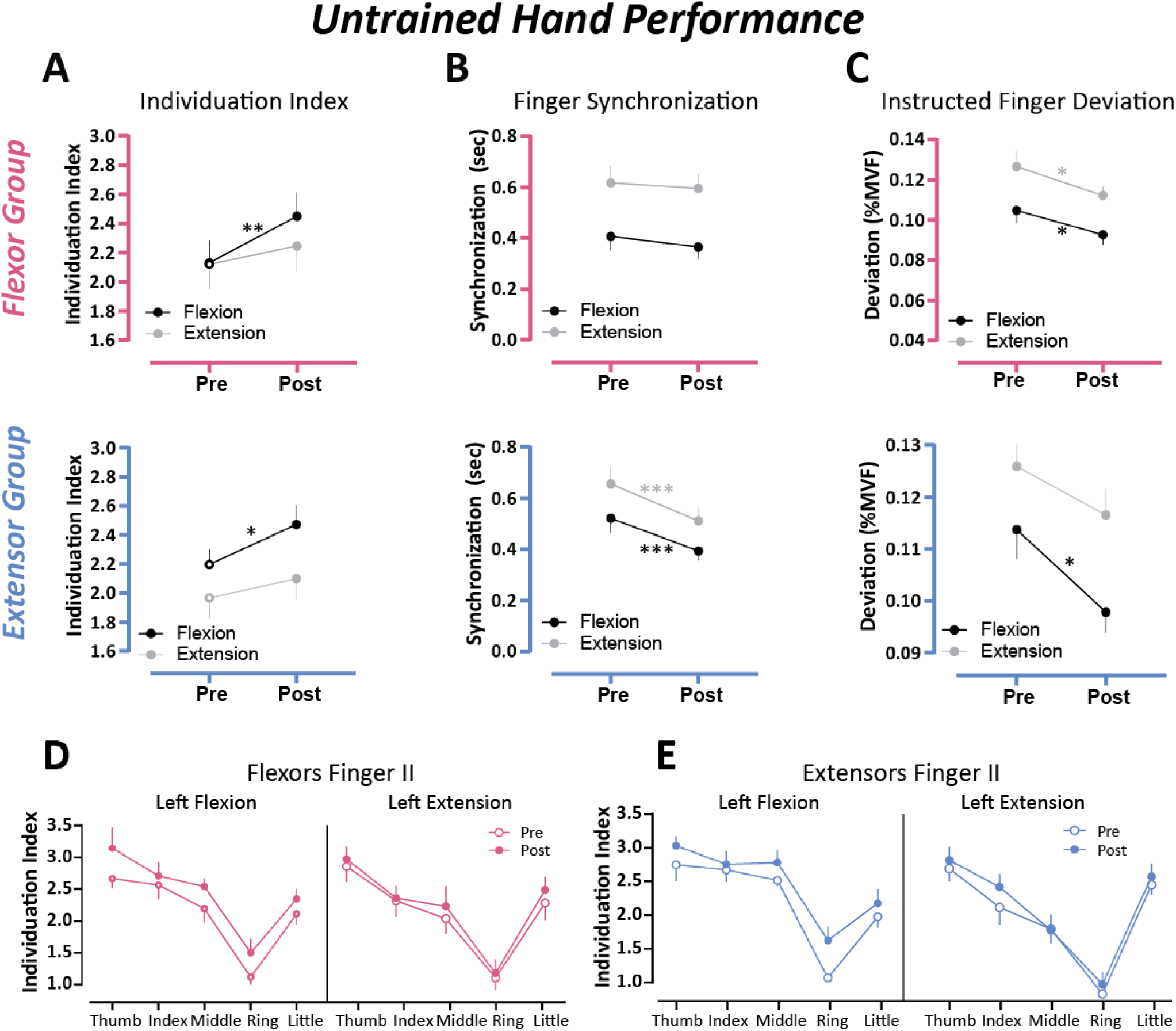
– Generalization of finger dexterity measures across hand, on left hand (untrained) movements for both Flexor and Extensor groups. (A) Training effect changes in Individuation Index. (B) Same as (A) but for changes in temporal synchronization between fingers. (C) Same as (A) but for force control, as calculated by MeanDevI, normalized by the MVF. (D) The Flexor group’s finger II values on LH flexion and LH extension, pre and post training. (E) The Extensor group’s finger II values on LH flexion and LH extension, pre and post training.

The average temporal synchronization of LH flexion of the flexor group prior to the training was 399.6 (±54.4) ms, and following training was 374.5 (±48.3) ms, resulting in an improvement of Δ=25.1 ms (Figure 5B, top panel). Two-way (time vs. direction) RM-ANOVA was performed for the flexor group, which showed a significant effect for the direction parameter (F(1,11)=25.85, p=0.0004), though not for the time parameter (F(1,11)=0.519, p=0.4863), nor an interaction between the two (F(1,11)=0.1413, p=0.7141). For the extensor group, the average LH extension synchronization (±SE) prior to training was 663.3 (±64.2) ms, and decreased to 518.0 (±54.9) ms post-training, resulting in an improvement of Δ=145.2 ms (Figure 5B, bottom panel). Two-way (time vs. direction) RM-ANOVA was also performed for the extensor group, which showed a significant effect for both the time parameter (F(1,11)=7.03, p=0.0225) and the direction parameter (F(1,11)=22.91, p=0.0006), though there was no interaction between the two (F(1,11)=0.1578, p=0.6988).

The average accuracy, measured as *meanDevI* normalized by MVF, of LH flexion of the flexor group prior to the training was 0.102 (±0.0056), and following the training was 0.0925 (±0.005), resulting in an improvement of Δ=0.0095 (Figure 5C, top panel). Two-way (time vs. direction) RM-ANOVA was performed for the flexor group, which showed a significant effect for both the time parameter (F(1,11)=11.28, p=0.0064) and the direction parameter (F(1,11)=57.54, p<0.0001), though there was no interaction between the two (F(1,11)=0.1439, p=0.7117). For the extensor group, the average LH extension accuracy (±SE) prior to training was 0.1275 (±0.0053), and decreased to 0.1329 (±0.0228) post-training, resulting in an improvement of Δ=0.0054 (Figure 5C, bottom panel). Two-way (time vs. direction) RM-ANOVA was also performed for the extensor group, which showed a significant effect for both the time parameter (F(1,11)=5.507, p=0.0369) and the direction parameter (F(1,11)=21.25, p=0.0006), as well as an interaction between the two (F(1,11)=10.67, p=0.0085).

The remaining movement type to investigate is the combination of both lateral and directional generalization, i.e. movement in the opposite hand and opposite direction of the trained movement. For the flexor group, this was the LH extension motion, for which pre-training they had an average II (±SE) of 2.12 (±0.17), and 2.25 (±0.18) post-training, resulting in an improvement of Δ=0.13 (Figure 5A, top panel). Post-hoc tw-tailed, paired sample t-test comparison between performance on Day 5 and Day 1 showed that training did not lead to significant improvement in left hand extension II values (t(12)=1.617, p=0.132). The extensor group, however, did have significant improvement in their LH flexion II. Pre-training, the extensor group’s average LH flexion II was 2.20 (±0.11), and 2.47 (±0.13) post-training, resulting in an improvement of Δ=0.27 (Figure 5A, bottom panel). Post-hoc two-tailed, paired sample t-test comparison between performances on Day 5 and Day 1 showed that training led to significant improvement in left hand flexion II values (t(12)=3.264, p=0.014). Figure 5D-F shows the individual fingers’ II values in both directions, and clearly shows the lack of improvement in LH extension in both groups, though a slight improvement across all fingers in LH flexion for both the flexor and extensor groups.

### Effects of prior experience on generalization

A cohort of musicians was used to examine the effect of prior experience on the ability to generalize motor learning. In order to enable proper comparison, the Musicians group completed the experiment in the same manner as the extensor group and trained only on RH extension motions. When looking at the performance during the training period, the daily averages (±STD) of the active deviation were 9.14 (±0.49)%, 7.83 (±0.43)%, and 7.43 (±0.22)% for days 2-4, respectivitly, with an improvement of 2.69% (i.e., 9.89%-7.20%) between the first and last blocks; the daily averages of the passive deviation were 2.81 (±0.46)%, 2.42 (±0.33)%, and 2.24 (±0.22)% for days 2-4, respectively, with an improvement of 1.53% (i.e., 3.52%-1.99%) between the first and last blocks. The daily averages of the reaction time were 505.9 (±51.7) ms, 416.8 (±29.4) ms, and 395.6 (±12.2) ms for days 2-4, respectively, with an improvement of 184.7 ms (i.e., 575.5-390.8 ms) between the first and last blocks. The daily averages of the synchronization were 364.4 (±1.2) ms, 268.9 (±14.3)ms, and 243.8 (±20.0)ms for days 2-4, respectively, with an improvement of 169.2ms (i.e., 363.7-194.5ms) between the first and last blocks. Overall, the values attained by the musician group were better than those of the extensor group (see Figure 6).

**Figure 6.**
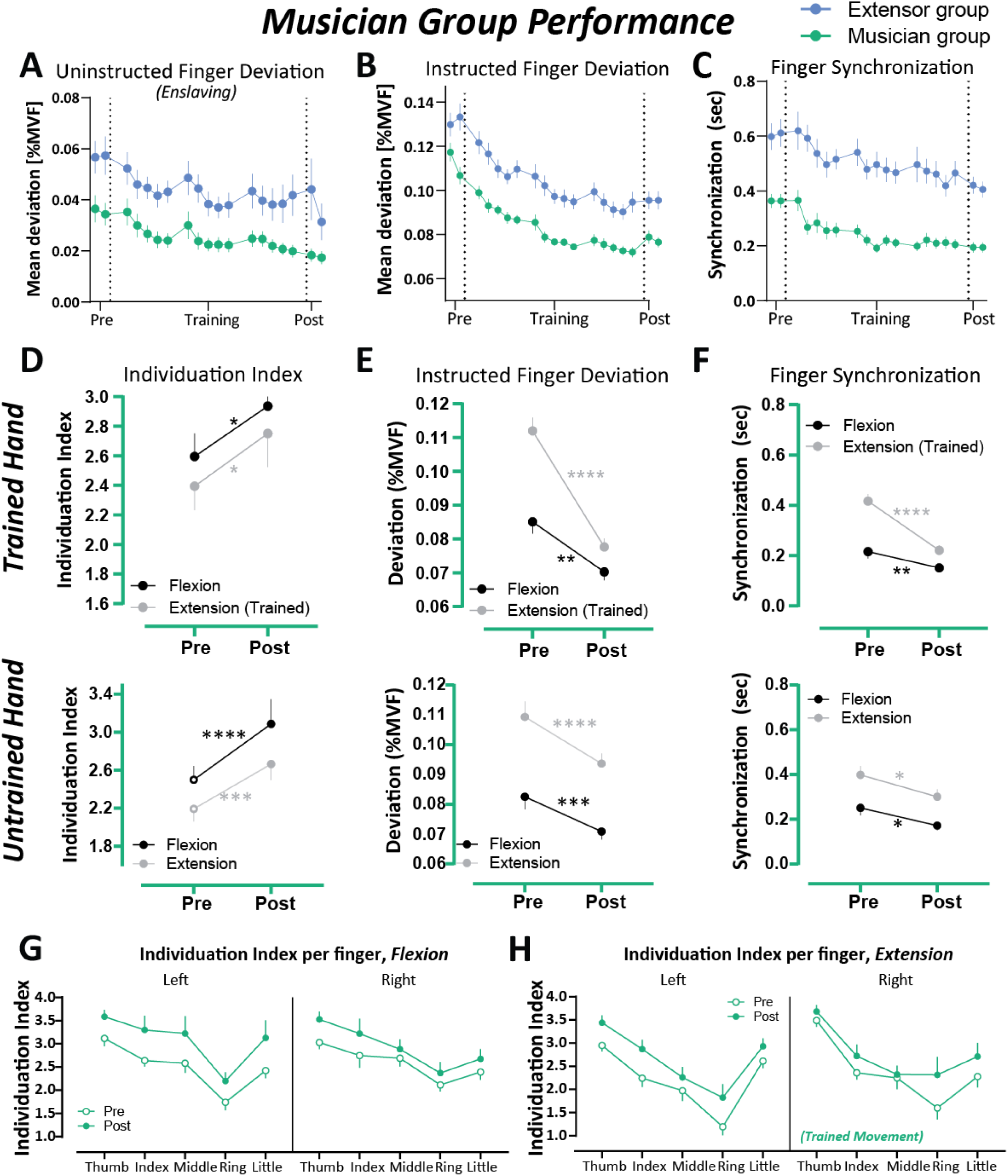
– History-dependent generalization of finger dexterity. Performance and training effect on the trained hand and ed direction of both musician and extensor groups. Training Performance metrics represent the average performance in of the 19 blocks; Pre and Post values are the average performance during the averaged values of the two testing blocks. The force control metric is depicted using the ‘Instructed Finger Deviation’, or MeanDevI, normalized by the MVF. (B) The structed Finger Deviation, or MeanDevU, normalized by the MVF. (C) The temporal synchronization between fingers. as n in the learning curves, as well as the training effect. (d-f) The training effect on the musician group. The top row esents RH motions and the bottom row represents LH motions. (D) Shows the force control, as calculated by MeanDevI. Shows the synchronization between fingers. (F) Shows the changes in Individuation Index. The musician group had ficant improvement in all metrics and in all motion types. (G) II values per finger on the flexion movements, both LH and flexion pre and post training. (H) II values per finger on the extension movements, both LH and RH flexion pre and post training.

#### Trained Direction Learning

In regards to the IIs that were measured on the first and last day of the study, the average RH extension II (±SE) of the musician group was 2.39 (±0.16) and 2.75 (±0.23), repectively, resulting in an average improvement of 0.36. Post-hoc two-tailed, paired sample t-test comparison between performance on Day 5 and Day 1 showed that training led to significant improvement in right hand extension II values (t(12)=2.660, p=0.010).

#### Opposite Direction Learning

For the musician group, the average RH finger flexion II (±SE) values on the first day for the thumb, index, middle, ring, and little fingers were 3.03 (±0.15), 2.75 (±0.26), 2.69 (±0.18), 2.12 (±0.14), and 2.39 (±0.17), respectively, with an overall hand average of 2.60 (±0.16). Following the training, the average RH flexion finger II values of the thumb, index, middle, ring, and little fingers for the musician group were 3.53 (±0.17), 3.22 (±0.32), 2.88 (±0.21), 2.37 (±0.24), and 2.68 (±0.21), respectively, with an overall hand average of 2.94 (±0.21). Post-hoc right-tailed, paired sample t-test comparison between performance on Day 5 and Day 1 showed that training led to significant improvement in right hand flexion II values in the musician group (t(12)=2.893, p=0.007). Figure 6G-H (right panels) shows the impact of each fingers’ II value on the overall hand II change. Though both directions improved across all fingers, improvements in RH flexion were more consistent than RH extension, where some fingers had a larger influence on the overall improvement. Two-way (time vs. direction) RM-ANOVA was performed and showed a significant effect for the time parameter (F(1,12)=12.47, p=0.0041), a significant, yet marginal, effect for the direction parameter (F(1,12)=4.617, p=0.0528), and no interaction between the two (F(1,12)=0.010, p=0.9224).

#### Opposite Hand Learning

For the musician group, the average LH finger extension II (±SE) values on the first day for the thumb, index, middle, ring, and little fingers were 2.95 (±0.13), 2.34 (±0.19), 1.97 (±0.22), 1.93 (±0.18),and 2.61 (±0.15), respectively, with an overall left hand average of 2.19 (±0.13). Following the training, the average LH finger extension II values of the thumb, index, middle, ring, and little fingers for the musician group were 3.44 (±0.16), 2.87 (±0.20), 2.26 (±0.22), 1.82 (±0.29), and little-2.93 (±0.17), respectively, with an overall left hand average of 2.66 (±0.17). Post-hoc right-tailed, paired sample t-test comparison between performance on Day 5 and Day 1 showed that training led to significant improvement in left hand extension II values (t(12)=3.843, p=0.001).

#### Opposite Hand and Opposite Direction Learning

For the musician group, the average LH finger flexion II (±SE) values on the first day for the thumb, index, middle, ring, and little fingers were 3.12 (±0.17), 2.64 (±0.13), 2.58 (±0.21), 1.74 (±0.17), and 2.42 (±0.16), respectively, with an overall left hand average of 2.50 (±0.14). Following the training, the average LH finger flexion II values of the musician group for the thumb, index, middle, ring, and little fingers were 3.59 (±0.15), 3.30 (±0.31), 3.22 (±0.37), 2.20 (±0.19), and 3.13 (±0.38), respectively, with an overall left hand average of 3.09 (±0.26). Post-hoc right-tailed, paired sample t-test comparison between performance on Day 5 and Day 1 showed that training led to significant improvement in left hand flexion II values (t(12)=3.743, p=0.001). Figure 6G-H (left panels) show the impact of each finger’s II value on the overall left hand II changes. Both directions improved across all fingers in a rather consistent manner. Two-way (time vs. direction) RM-ANOVA was performed for LH flexion and extension, which showed a significant effect for both time (F(1,12)=17.42, p=0.0013) and direction parameters (F(1,12)=8.81, p=0.0117), though no interaction between the two (F(1,12)=0.894, p=0.3631).

### Control experiment

To rule out the possibility that the observed improvement in dexterity is due to re-exposure to the test itself and not generalization of what has been learned, we recruited a control group consisting of naive participants (n=8) which performed the baseline testing session on Day 1, rested for 3 days and then were re-exposed to the same testing session on Day 5 (Supplementary Figure 5). We found that none of the dexterity measures showed significant improvement on Day 5 (statistical tests in all measures revealed non-significant time effect, *p* > 0.149, nor time×direction interaction effect, *p* > 0.153). Specifically, average RH II for flexion on the first day was 2.51 (±0.16), and on the last day was 2.63 (±0.09), resulting in a change of 0.12; average RH II for extension on the first day was 2.37 (±0.21), and on the last day was 2.41 (±0.14), resulting in a change of only 0.04; average LH II for flexion on the first day was 2.81 (±0.18), and on the last day was 2.74 (±0.09), resulting in a decrease of 0.07; average LH II for extension on the first day was 2.46 (±0.15), and on the last day was 2.47 (±0.06), resulting in a change of only 0.01. In terms of finger synchronization, the changes in finger synchronization for RH flexion, RH extension, LH flexion, and LH extension were -39 ms, -74 ms, 3 ms, and 12 ms, respectively. The changes in instructed finger deviation for RH flexion, RH extension, LH flexion, and LH extension were -1.7% MVF, -0.8% MVF, 0% MVF, and 0% MVF, respectively. The changes in uninstructed finger deviation for RH flexion, RH extension, LH flexion, and LH extension were -0.4% MVF, -1.4% MVF, 0.1% MVF, and -0.3% MVF, respectively. The lack of significant change in performance supports that the training is the main cause for the improvement seen in the flexor, extensor and musician groups. Moreover, this data strongly suggests that the asymmetric and history-dependent effects we observed in the main experiment is attributed to the generalization of the learning and not to re-exposure to the test for the second time, nor to the passage of time.

## Discussion

Our study provides detailed characterization of dexterous single and multi-finger flexion and extension movements in human participants. Dexterous ability was decomposed into components (finger individuation, force control, and temporal synchronization) which were investigated individually and collectively to shed light on the motor learning abilities during the training period, as well as the overall changes after the training period. Our goal was to further define the control mechanisms affecting dexterous movement, specifically the learning and generalization of flexion and extension in human multi-finger movements. Lastly, we investigated whether the learning and generalization of multi-finger dexterous movements were affected by the history of prior actions.

### Training effect on multi-finger dexterity components

In order to illuminate potential relationships of the neural control mechanisms, two cohorts of naïve participants were recruited and trained in either flexion or extension movements. The assignment of participants to the flexor or extensor group was done randomly, as is attested to by the similar baseline values of the groups. Differing values between the groups can be seen within the first blocks of training, especially in the accuracy and synchronization metrics (**Figure 3**). Baseline single and multi-finger actions revealed increased accuracy and enhanced synchronization in the flexion direction compared to the extension direction. This can be attributed in part to the fact that producing an extension motion is less common in human daily functions and therefore will inherently be performed in a less skilled manner^19^. This is also consistent with the general behavioral tendency evident in the design of human tools, such as the keyboard, piano, which rely on finger flexion, rather than extension when precise finger control is needed^28^. When participants were required to produce maximal contractions in the flexion and extension direction, Yu et al., (2010)^19^ showed that there is a higher force deficit (less accurate) in extension than that in flexion movements.

When a subject intends to exert force with one or more digits, the tendency of additional digits to unintentionally exert lesser forces has been described most often as “enslaving,” i.e., force production by the master digit(s) enslaves unintended force production in other digits^16, 29, 30^. Enslaving can be viewed as the inverse of individuation, in that a lower degree of enslaving corresponds to a higher degree of finger individuation. Our baseline data, however, showed no significant differences in enslaving between the extension and flexion direction. This result somehow contradicts previous reports showing that enslaving is higher during production of extension forces compared to that in flexion forces^19, 31^. One potential explanation of the lack of differences in enslaving between flexion and extension at baseline might be related to hand orientation while performing the task. While in most previous studies that reported increased enslaving in extension the orientation of the hand was in the prone position, in our setup the hand was placed in the neutral position. Previous work showed that hand posture has a significant effect on the excitability of corticospinal output^32^. Studies which used transcranial magnetic simulation (TMS) protocols to measure intracortical inhibition of M1^33−^

^36^ during a pinch precision task in different hand postures found that cortical inhibition decreased during hand pronation and supination compared to a neutral posture. Although a direct examination of the contribution of the intracortical inhibition phenomena to finger individuation ability in humans remains unknown, we speculate that the reduced inhibition in the pronation orientation might be related to the enslaving of the uninstructed fingers. Another explanation of the similar baseline enslaving values of flexion and extension motions might be the fact that our participants produced much lower forces in the instructed fingers compared to previous work during the individuation task, potentially due to the altered orientation. Indeed, enslaving of non-instructed digits increases as the master digit produces more force^15^. Whether the origin of the differences between the extension and flexion direction is due to difference in neural control, biomechanical coupling (connections between tendons and muscles) and/or in the structure of motor units of the corresponding muscles, is yet to be determined.

The learning curves during the three days of training (**Figure 3**) show that both groups had trends of improvement, though the flexor group consistently performed better across the different dexterity metrics. The improvements were evident between the five individual blocks within a single day of training, as well as over the three training days, and can be seen clearly in **Figure 3**. By decomposing the movement into the three components (i.e., finger individuation, force control and temporal synchronization), the differences in their contribution to the overall improvement during the training were unveiled. Though all three dexterity parameters improved during the training period, the element which improved most significantly was the force control and the ability to obtain the target force more accurately.

When looking at the temporal realm, both the reaction time and the multi-finger synchronization improved following training. Improvement in finger dexterity measures within multiday training in multi-finger motor skill (termed “synergy”) was seen previously^37^, yet it was specific to the flexion direction. In this study, multiday training led to reduced deviation (e.g., combined metric of force control of the instructed finger and enslaving of the uninstructed fingers), faster RT responses and rapid execution times in the flexion direction (the only tested direction). Interestingly, Waters et al., 2016^37^ showed that these dexterity components were more augmented if bihemispheric anodal transcranial direct current stimulation (tDCS) was applied to the primary motor cortex (M1) during the training, suggesting for potential involvement of the motor cortex in the control process of some aspects of finger dexterity. A study which trained participants in a subset of multi-finger (2 and 3 fingers) chords in the flexion direction found significant generalization in the reaction time and accuracy for chords composed of novel configurations of the practiced elements (i.e., fingers), and chords that contained a new element^38^. Our study goes beyond these works by showing that despite inferior dexterity in the extension direction at baseline, multiday extension-based training improved individuation, force control and synchronization in the trained direction. Unfortunately, our results cannot reveal whether similar or separate neural mechanisms underlie the control of individuation, force control and temporal synchronization during multi-finger flexion and extension movements. Our lack of correlation results (Supplementary Figure 6) between the changes of the dexterity components following training, however, indicates that these variables might be derived from dissociable mechanisms. In addition, we confirm that the reported results are attributed to the generalization effect and not to re-exposure to the test for the second time, or due to passage of time between the baseline and post-training evaluation tests.

### Direction-dependent generalization of finger dexterity

After quantifying finger dexterity components and characterizing the learning processes of each component following multi-day training, our second primary aim was to evaluate generalization of the improved skill between the flexion and extension direction within the trained hand. The flexor group trained on a multi-finger chord task that required quick, synchronized production of difficult finger flexor muscle activation patterns whereas the extensor group used the same training regimen though using finger extensor muscles.

It is known that the context in which an individual trains affects whether learning generalizes or transfers to untrained movements^3, 39, 40^. Here, the direction of movements (for example, extension) could be considered the ‘context’ in which learning occurs. Both finger flexion and extension movements are strictly context dependent as each engage separate groups of muscles with distinct activity patterns. Thus, testing the opposite and untrained direction is considered testing the generalization to a novel context. Although flexion and extension naturally co-occur in most hand functions, and the success in performing these tasks depends on the ability to precisely coactivate finger flexor and extensor muscles in both hands, we found that transfer across contexts was direction specific with the extension movement affecting the flexion movement, but lack of clear generalization from flexion to extension movement. The more pronounced asymmetry between the two groups can also be seen in the changes in the Individuation Index before and after training. Though both groups had significant improvements in their trained hand and direction, only the extensor group had significant improvement in the opposite direction on the same hand.

The asymmetrical generalization pattern supports the *biased-overlap hypothesis* when considering the control hypotheses introduced earlier. But why is generalization of finger movements biased toward a specific direction? We interpret this direction specificity, or asymmetry, as a reflection of the differences in neural substrate of the underlying flexion versus extension dexterity. Previous work has shown that pre-existing bias of flexion is evident by the ability to more precisely control movements of finger flexors compared to extensors^6^. Additionally, clinical data for stroke survivors with cortical lesions who, despite regaining good flexion-based grasp, have very weak finger and wrist extensors, preventing hand opening^7–12^. Lastly, a recent neurophysiological study showed increased representation of finger flexion, but weaker representation of finger extension, using micro-stimulation of the human motor cortex^13^. Nevertheless, interpretation of this result is complicated and should be treated carefully because a change in direction not only changes the context, but also the force level needed during flexion vs. extension. Thus, it cannot be concluded that the asymmetrical generalization effect is purely a contextual direction effect.

### Lateral, across-hand, generalization of finger dexterity

This pattern of generalization is not limited to the across-direction of the trained hand but is also true when individuals generalized what has been learned to the untrained hand. The flexor group had significant improvement in the trained direction in the opposite hand, whereas the extensor group did not. It is interesting to note, however, that the extensor group improved in untrained finger flexion, which could theoretically be the lateral reflection of the directional generalization discussed earlier. The flexor group, which did not improve in trained finger extension, also did not demonstrate improvement in untrained finger extension. Neither the flexor nor extensor group obtained significant improvement in untrained extension.

Flexion-specific generalization to the untrained hand was previously reported in healthy naïve participants^37, 38, 41^ and stroke patients^26^. For example, we previously found that intensive training of the paretic hand with a multi-finger task in chronic ischemic stroke patients for 5 days improved finger individuation, not only in the trained hand, but also generalized to the untrained non-paretic hand, and lasted for at least 6 months following the training^26^. Consistently, other aspects of finger dexterity including force accuracy and reaction times were also generalized to the flexion direction of the untrained hand. This effect was further enhanced if training was coupled with bihemespheric tDCS M1, suggesting effector-independent representations of dexterity that allowed, in part, intermanual transfer that benefitted both hands^37^. One explanation of the generalization to the flexion direction of the untrained hand might be attributed to the role of the ipsilateral hemisphere during unimanual hand training. Previous neuroimaging studies have consistently observed ipsilateral activation in the sensorimotor cortex. Many of these studies used a sequential finger opposition movement and reported ipsilateral activation^42, 43^, while others used complex and sequential movements and reported even larger increased activation in ipsilateral parietal and premotor regions ^38, 44^. It is possible to think that the ipsilateral hemisphere may contribute to rapid selection of learned sensorimotor associations, and this involvement is especially employed during complex actions, such as in the case of our multi-finger dexterity training^45, 46^. In a series of experimental conditions, Hazeltine et al., (2007)^38^ conveniently showed that the strong ipsilateral activity in the left motor cortex was specific to the execution of complex movements independent of the sequential nature of the task.

While this interpretation might explain the generalization effect we observed in the flexion direction of the untrained hand, it cannot explain why participants failed to generalize the improved dexterity to the extension direction of the untrained hand. The complexity and task demands were similar across directions but generalization to the untrained hand was specific to the flexion direction only. One explanation for the direction-specific generalization across hands might be related to the fact that ipsilateral activation during unimanual movement might be sensitive not only to the task complexity demand, but also to other high-level parameters such as movement direction and/or amplitude^47–49^, and that this direction-sensitivity is biased more toward the flexion direction. An alternative explanation for the lack of generalization of extension movement might be related to the fact that these movements are less used in our daily function and therefore are less represented in the motor cortex. The structure of activation patterns is determined by the way we use our hands in everyday life^50, 51^. Using fMRI paradigms, Ejaz et al (2015)^51^ found that frequently co-occurring finger movements in the flexion direction led to strong associations between the cortical modules that encode them. Since extension-based movement is less frequent in our daily function, we speculate that the cortical association of these movement might be less represented compared to the flexion direction. In addition, using direct electrical stimulation over the human motor cortex, flexion of the fingers was evoked far more often than extension (flexion 63 times, extension 8)^6^. This finding was further discussed and interpreted by as the fact that the representation of finger flexion is more ‘magnified’ than that of extension because almost all functions of the hand in grasping objects and using tools require strong and/or precise finger flexion, with extension being used simply to release or withdraw the fingers^52^.

The hand use interpretation of generalization makes an interesting prediction: the amount of generalization is influenced by the prior history of practiced action. Consequently, if a group of participants had frequently used their finger extensors during daily life, they should show increased generalization also for the untrained hand in the extension direction. We tested this idea by recruiting musicians (pianists and violinists) with years of practice in both the extension and flexion direction and examined the generalization patterns across untrained direction and hand.

### History-dependent learning and generalization of finger dexterity

The effect of prior experience with dexterous movement training was examined by recruiting a third cohort comprised of experienced musicians who performed the study in an identical manner to the extensor group. All baseline dexterity parameters were pronouncedly superior in the musician group, though both groups improved during the training period. Similar to the extensor group, the musician group exhibited directional generalization in the untrained flexion direction of the trained hand, as well as intermanual transfer (across hands) in the untrained hand flexion. Though both groups improved, the learning in the musician group was more pronounced than in the extensor group. Interestingly, the musicians presented lateral generalization with an improvement in untrained extension direction of the untrained hand, whereas the extensor group did not. The musician group was the only group in the study to significantly improve in all metrics and in all directions. These results are indicative of modular and rather strong representation of both flexion and extension movement in musicians, confirming the prediction of prior history effect on generalization of multi-finger dexterous movements. Indeed, intensive finger training in musicians not only affects behavior ^21, 53^, but also elicits experience-dependent changes in the modular architecture of M1^54–56^, and this is in addition to the fact that the cortical representation is affected by everyday hand use^51^.

### Limitations

An additional possible explanation for the generalization effect we observed following the learning may be related to the perceptual representations alongside the motor representations. In terms of perceptual learning, participants might become more skilled at recognizing the complex patterns formed by the arrays of force targets that define the stimuli patterns^57–59^, regardless of the movement direction. In terms of motor learning, participants might become more adept in producing particular multi-finger chord responses with their hands. It is also possible that training leads to both perceptual and motor improvements, or improvements at an intermediate level in which specific input patterns are associated with particular chord responses. Although perceptual learning was not tested directly in our experiments, the perceptual stimulus-response hypothesis is unlikely to account for the asymmetrical generalization we observed, since both extension and flexion actions required responses to similar stimuli, yet the motor behavior was different. In agreement with the response-based hypothesis, previous work showed that participants were faster in responding to novel stimuli that mapped to practiced multi-finger chord responses compared to novel stimuli that mapped to unpracticed chord responses, suggesting that learning in the chord is largely response based^38^.

In addition, our study was limited by mechanical and technical aspects of the ergonomic measurement device, specifically maximal hand size and maximal force capability. Nevertheless, we do not expect these limits to affect the validity of the forces measured, nor the recruitment of a wide variety of population. Additionally, some cognitive processes, such as familiarization with the setup and developing strategies during learning the task, may have contributed to the improvement in dexterity we detected. Under the assumption that these cognitive processes equally affect the performance regardless of the type and direction of action, they should not affect the asymmetry presented in the improvements of the different direction. Future studies are needed to precisely evaluate the contribution of the cognitive functions during flexion/extension finger dexterity.

### Summary and Conclusions

The evidence that generalization across directions can occur following uni-directional training supports the assumption that the control mechanisms of flexion and extension motions are intrinsically connected. When considering this as well as the cortical organization of flexor and extensor motor neurons from previous work, the *independent control hypothesis* proposed earlier does not seem like an appropriate control framework of finger dexterity. One possible, yet informative and direct, way to gain insight as to the proper functioning of a control system is to monitor its behavior when specific parts of it are affected due to illness or injury. When pertaining to the brain, this can be investigated in animal models or in patients with neurological injury, such as stroke. The asymmetry between flexion and extension control and abilities has been described in numerous studies concerning stroke survivors^10, 11, 26^, as well as in animal models with induced brain legions^9^. The results of the study further corroborate the stark asymmetry between improvements following flexion training and extension training, suggesting a directional dependency that is biased towards improvement in flexion motion. Therefore, the proposed flexor *biased-overlap hypothesis* (Fig. 1D, most right panel) appears to adequately reflect the control mechanism governing single- and multi-finger dexterous movement.

The current study characterizes the behavioral principles underlying the control process, learning and generalization of dexterous extension and flexion movements in a population of healthy young individuals. Our data indicate that control of multi-digit dexterous patterns is direction-specific in humans, supporting the biased-overlap hypothesis by which the control circuits for learning of finger flexion and extension are overlapped (i.e., they partially, but asymmetrically, transfer between directions and hands). Moreover, participants with extensive prior history in dexterous training exhibited increased improvement compared to their counterparts. These findings can be utilized in future studies, as well as in clinical settings such as rehabilitation protocols for those with unilateral or unidirectional hemiparesis stemming from neural impairments.

## Acknowledgements

We would like to thank Rotem Lootsky for his help in designing and 3D printing the MRI-compatible dexterity device, and Jeries Saleh for his assistance in running the experiment in the musician group. This study was funded by the Israel Science Foundation (ISF) Grant 1634/19 (FM) and German-Israel Foundation (GIF) Grant I-2535-409.10/2019 (FM).

## Conflict of Interest

The authors declare no competing financial interests.

## Supplemental Information

### Supplementary Tables

**Table S1.** – Demographic information of the three groups, Data (±STD)

|  |  | Flexors | Extensors | Musicians | Control |
| --- | --- | --- | --- | --- | --- |
| Participants | n | 13 | 13 | 13 | 8 |
| | Age | 26.9 ($\pm 3.5$) years | 26.2 ($\pm 4.9$) years | 24.2 ($\pm 3.7$) years | 23.3 ($\pm 1.7$) years |
| | Handedness (Edinburgh) | 90.0 ($\pm 8.7$) | 87.9 ($\pm 22.5$) | 76.7 ($\pm 22.8$) | 82.2 ($\pm 14.3$) |

**Table S2.** – Musical experience of the Musician group, according to instrument type

| Main Instrument | n | Age | Years Active |
| --- | --- | --- | --- |
| Piano | 7 | 24.9 ($\pm 3.1$) | 10.7 ($\pm 5.1$) |
| Violin/Viola | 5 | 23.2 ($\pm 2.2$) | 11.8 ($\pm 4.1$) |
| French Horn | 1 | 24 | 15 |
| **Total** | **13** | **24.2 ($\pm 2.7$)** | **11.5 ($\pm 4.5$)** |

**Table S3.** – The various Individuation Index values of the Flexor and Extensor groups are listed. The t-test used above is paired sample t-test with the Holm-Šídák correction for repeated measures.

| Parameter |  | pre<br>mean (SE) | post<br>mean (SE) | t<sub>(12)</sub> | p | pre<br>mean (SE) | post<br>mean (SE) | t<sub>(12)</sub> | p |
| --- | --- | --- | --- | --- | --- | --- | --- | --- | --- |
| Trained Direction,<br>Trained Hand // |  | <u>Right Hand Flexion</u> |  |  |  | <u>Right Hand Extension</u> |  |  |  |
|  | Mean | 2.36 (0.16) | 2.65 (0.17) | 3.717 | 0.003** | 2.25 (0.17) | 2.58 (0.14) | 3.006 | 0.011* |
|  | Thumb | 2.97 (0.12) | 3.12 (0.13) |  |  | 2.95 (0.21) | 3.40 (0.16) |  |  |
|  | Index | 2.67 (0.23) | 3.03 (0.25) |  |  | 2.55 (0.27) | 2.66 (0.19) |  |  |
|  | Middle | 2.37 (0.17) | 2.72 (0.21) |  |  | 2.09 (0.24) | 2.76 (0.22) |  |  |
|  | Ring | 1.66 (0.22) | 1.93 (0.20) |  |  | 1.17 (0.19) | 1.63 (0.24) |  |  |
|  | Little | 2.13 (0.17) | 2.44 (0.18) |  |  | 2.50 (0.29) | 2.44 (0.19) |  |  |
| Untrained Direction,<br>Trained Hand // |  | <u>Right Hand Extension</u> |  |  |  | <u>Right Hand Flexion</u> |  |  |  |
|  | Mean | 2.16 (0.14) | 2.31 (0.13) | 0.642 | 0.782 | 2.01 (0.18) | 2.62 (0.17) | 6.127 | 0.0001*** |
|  | Thumb | 2.94 (0.25) | 3.24 (0.14) |  |  | 2.63 (0.21) | 3.17 (0.18) |  |  |
|  | Index | 2.25 (0.22) | 2.37 (0.16) |  |  | 2.36 (0.29) | 3.14 (0.32) |  |  |
|  | Middle | 1.98 (0.19) | 2.07 (0.27) |  |  | 1.95 (0.28) | 2.64 (0.20) |  |  |
|  | Ring | 1.48 (0.23) | 1.53 (0.25) |  |  | 1.37 (0.19) | 1.99 (0.20) |  |  |
|  | Little | 2.16 (0.17) | 2.33 (0.17) |  |  | 1.73 (0.14) | 2.17 (0.12) |  |  |
| Trained Direction,<br>Untrained Hand // |  | <u>Left Hand Flexion</u> |  |  |  | <u>Left Hand Extension</u> |  |  |  |
|  | Mean | 2.13 (0.15) | 2.45 (0.16) | 4.103 | 0.003** | 1.97 (0.14) | 2.10 (0.14) | 1.556 | 0.146 |
|  | Thumb | 2.67 (0.15) | 3.14 (0.33) |  |  | 2.67 (0.19) | 2.80 (0.20) |  |  |
|  | Index | 2.56 (0.22) | 2.71 (0.21) |  |  | 2.10 (0.26) | 2.40 (0.20) |  |  |
|  | Middle | 2.20 (0.22) | 2.54 (0.12) |  |  | 1.79 (0.21) | 1.77 (0.23) |  |  |
|  | Ring | 1.12 (0.25) | 1.50 (0.22) |  |  | 0.83 (0.13) | 0.97 (0.18) |  |  |
|  | Little | 2.11 (0.17) | 2.35 (0.16) |  |  | 2.43 (0.15) | 2.55 (0.19) |  |  |
| Untrained Direction,<br>Untrained Hand // |  | <u>Left Hand Extension</u> |  |  |  | <u>Left Hand Flexion</u> |  |  |  |
|  | Mean | 2.12 (0.17) | 2.25 (0.18) | 1.617 | 0.132 | 2.20 (0.11) | 2.47 (0.13) | 3.264 | 0.014 * |
|  | Thumb | 2.84 (0.24) | 2.97 (0.20) |  |  | 2.75 (0.24) | 3.03 (0.14) |  |  |
|  | Index | 2.32 (0.25) | 2.36 (0.20) |  |  | 2.67 (0.18) | 2.75 (0.19) |  |  |
|  | Middle | 2.04 (0.24) | 2.23 (0.31) |  |  | 2.52 (0.09) | 2.78 (0.19) |  |  |
|  | Ring | 1.10 (0.19) | 1.18 (0.22) |  |  | 1.07 (0.23) | 1.63 (0.20) |  |  |
|  | Little | 2.28 (0.27) | 2.49 (0.21) |  |  | 1.98 (0.16) | 2.18 (0.21) |  |  |

### Supplementary Figures

**Supplementary Figure 1.**
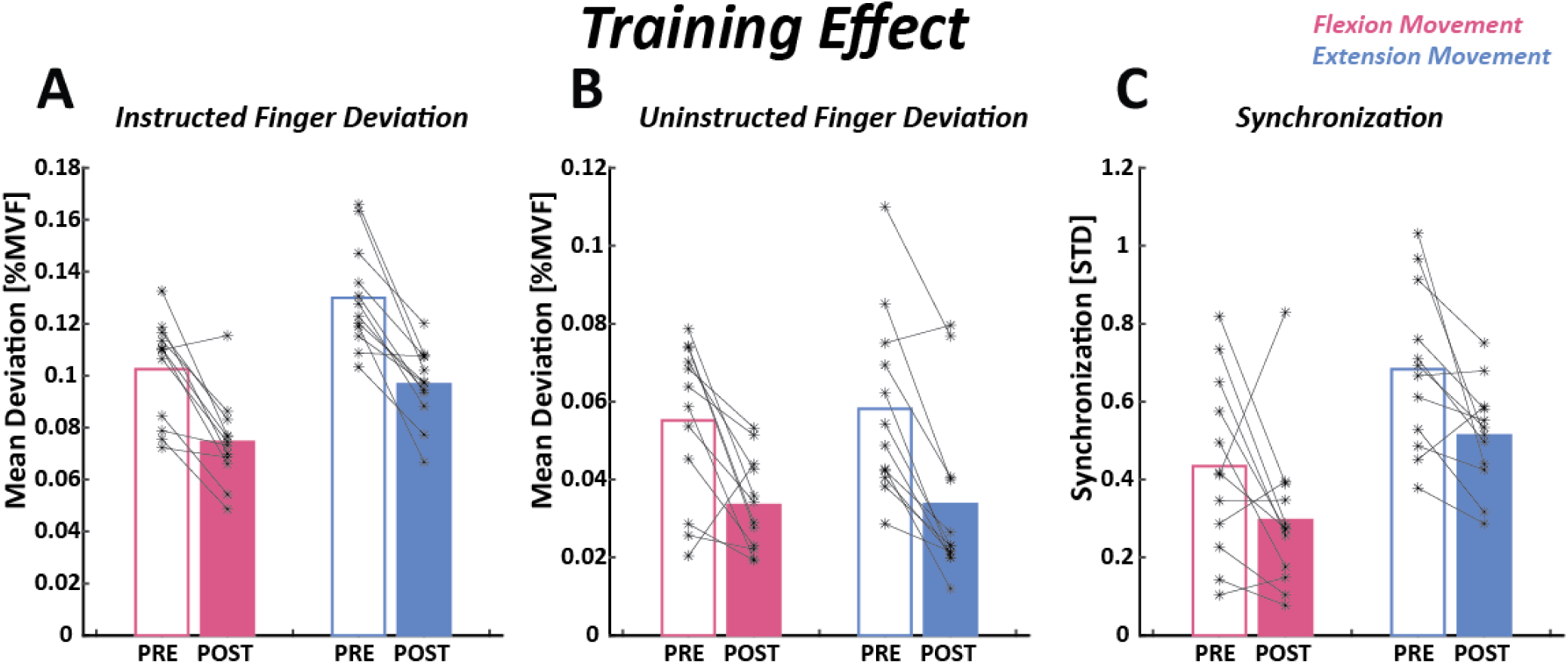
– Individual data of the training effect on both the flexor (pink) and extensor (blue) groups. Each participant is represented by two connected asterisks which show the change in performance between the Pre and Post days. The data represents performance during flexion motion for the flexor group, and extension motion for the extensor group. (A) The instructed finger deviation, normalized by the MVF, shows trends of improvements among most participants of both groups, regardless of baseline values.(B) The uninstructed finger deviation, normalized by MVF, also shows trends of improvement among most participants of both groups. (C) The synchronization improved in both groups in most participants of both groups.

**Supplementary Figure 2.**
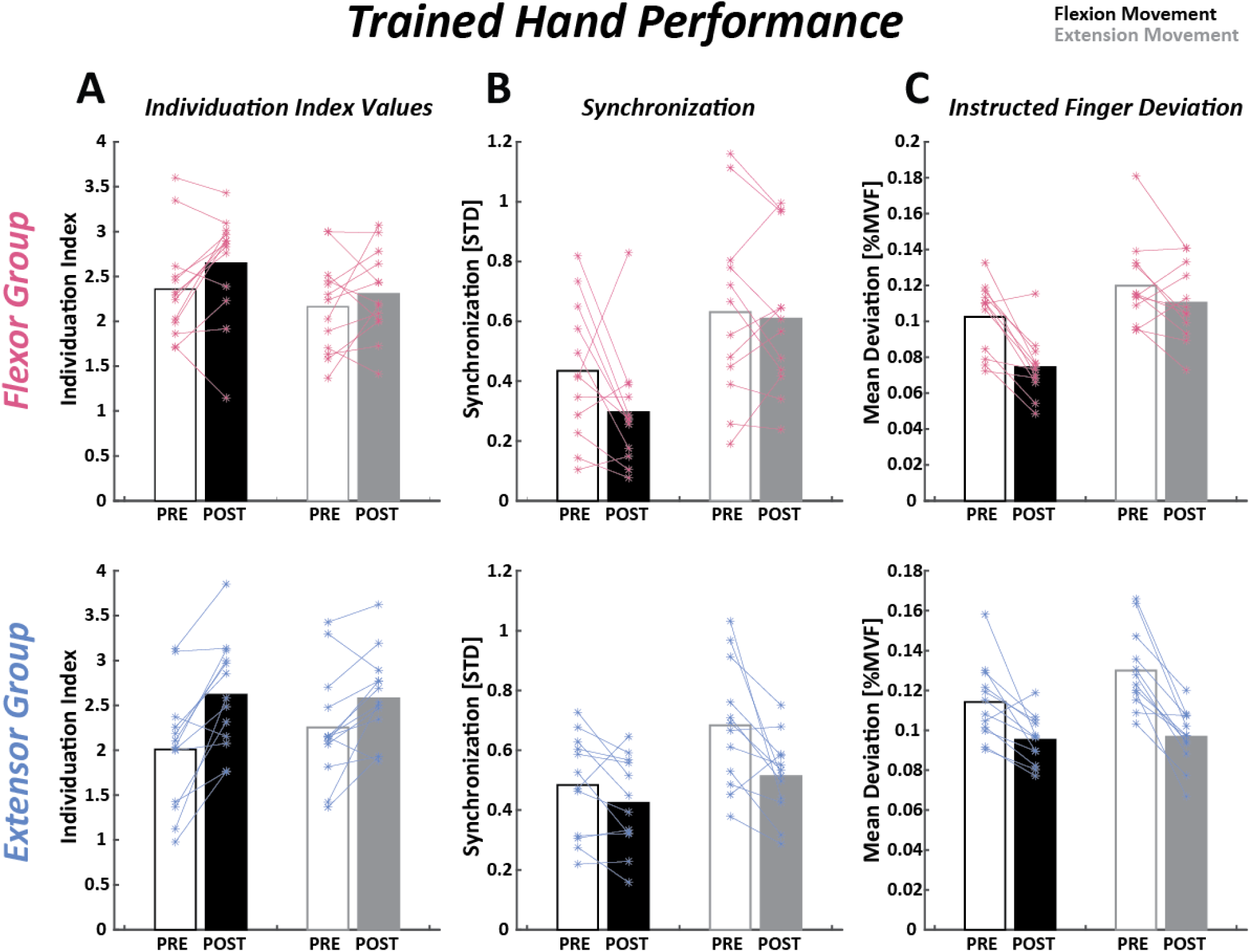
– Individual data of the flexor and extensor groups’ performance on the trained hand. Each participant is represented by two connected asterisks which show the change in performance between the Pre and Post days. Flexion motion is depicted in black and extension motion is depicted in grey. (A) The individuation index of the individual participants can be seen for both directions in both groups. (B) The synchronization of the individual participants can be seen for both directions in both groups. (C) The instructed finger deviation, normalized by the MVF, of the individual participants can be seen for both directions in both groups.

**Supplementary Figure 3.**
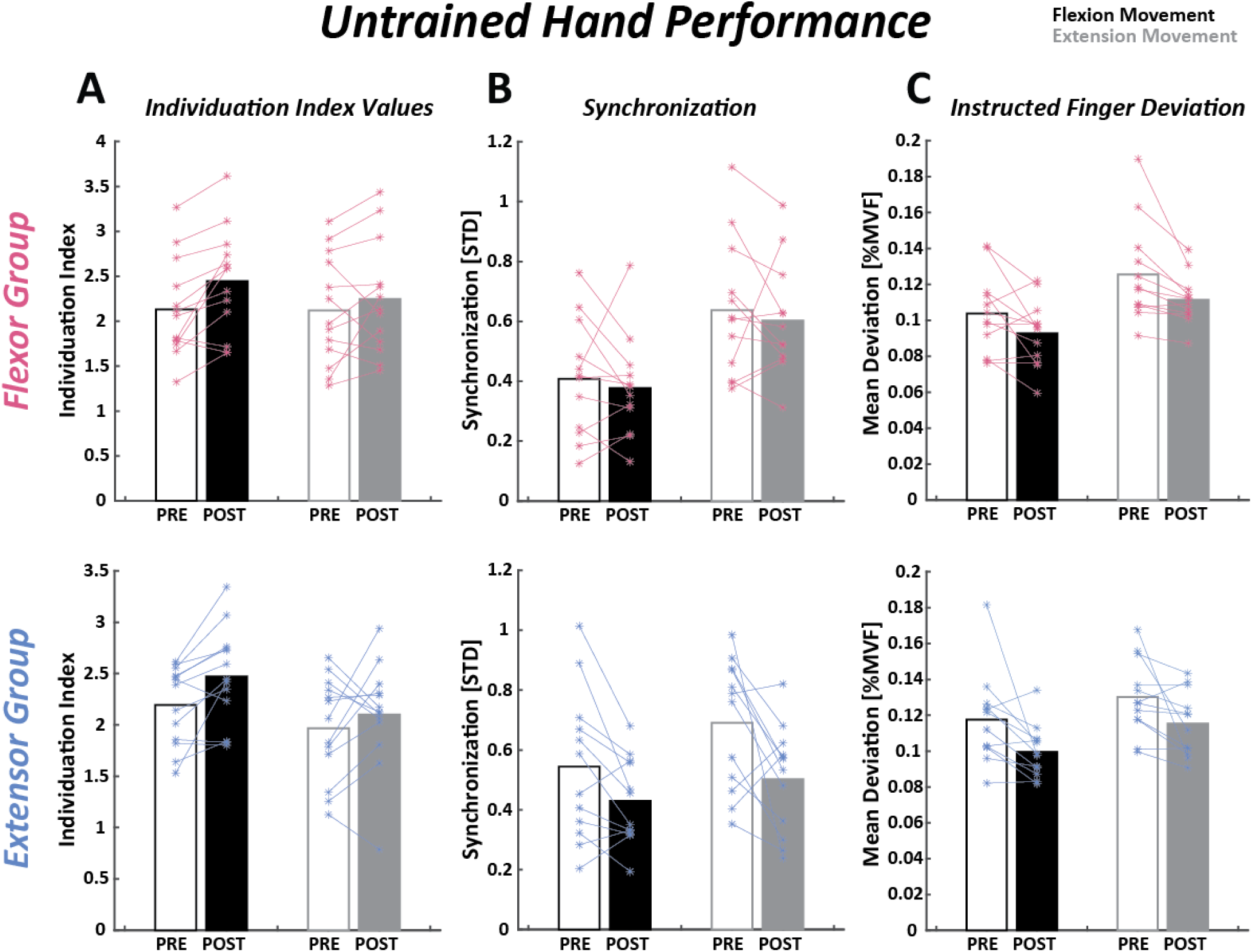
– Individual data of the flexor and extensor groups’ performance on the untrained hand. Each participant is represented by two connected asterisks which show the change in performance between the Pre and Post days. Flexion motion is depicted in black and extension motion is depicted in grey. (A) The individuation index of the individual participants can be seen for both directions in both groups. (B) The synchronization of the individual participants can be seen for both directions in both groups. (C) The instructed finger deviation, normalized by the MVF, of the individual participants can be seen for both directions in both groups.

**Supplementary Figure 4.**
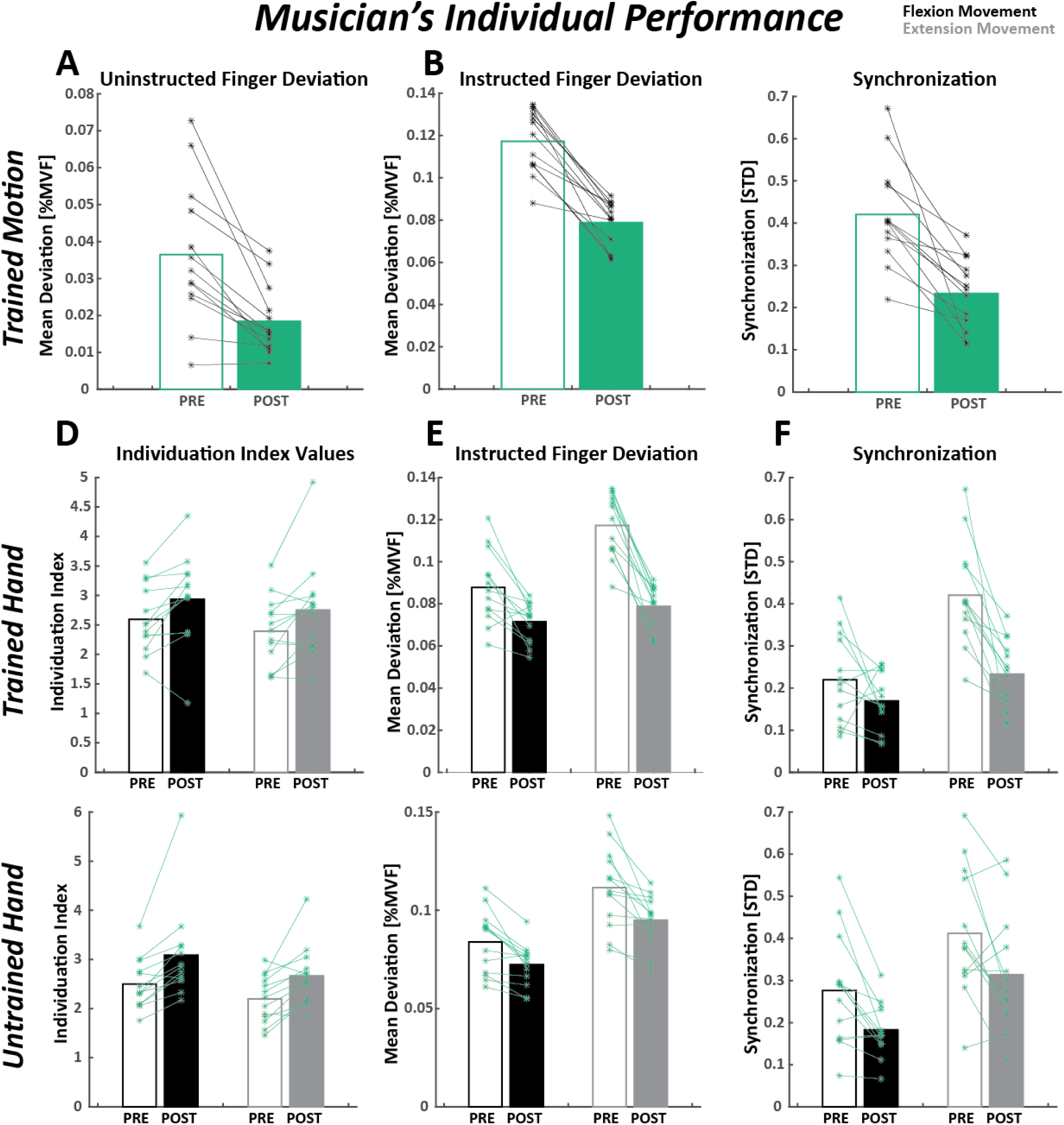
– Individual data of the musician group’s performance. Each participant is represented by two connected asterisks which show the change in performance between the Pre and Post days. Flexion motion is depicted in black and extension motion is depicted in grey. (A) Uninstructed finger deviation, normalized by MVF, of the individual participants for right hand extension motion. (B) Instructed finger deviation, normalized by MVF, of the individual participants for right hand extension motion. (C) Synchronization of the individual participants for right hand extension motion. (D) The individuation index of the individual participants can be seen for both directions for right hand (top) and left hand (bottom) movements. (E) The synchronization of the individual participants can be seen for both directions for right hand (top) and left hand (bottom) movements. (F) The instructed finger deviation, normalized by the MVF, of the individual participants can be seen for both directions for right hand (top) and left hand (bottom) movements.

**Supplementary Figure 5.**
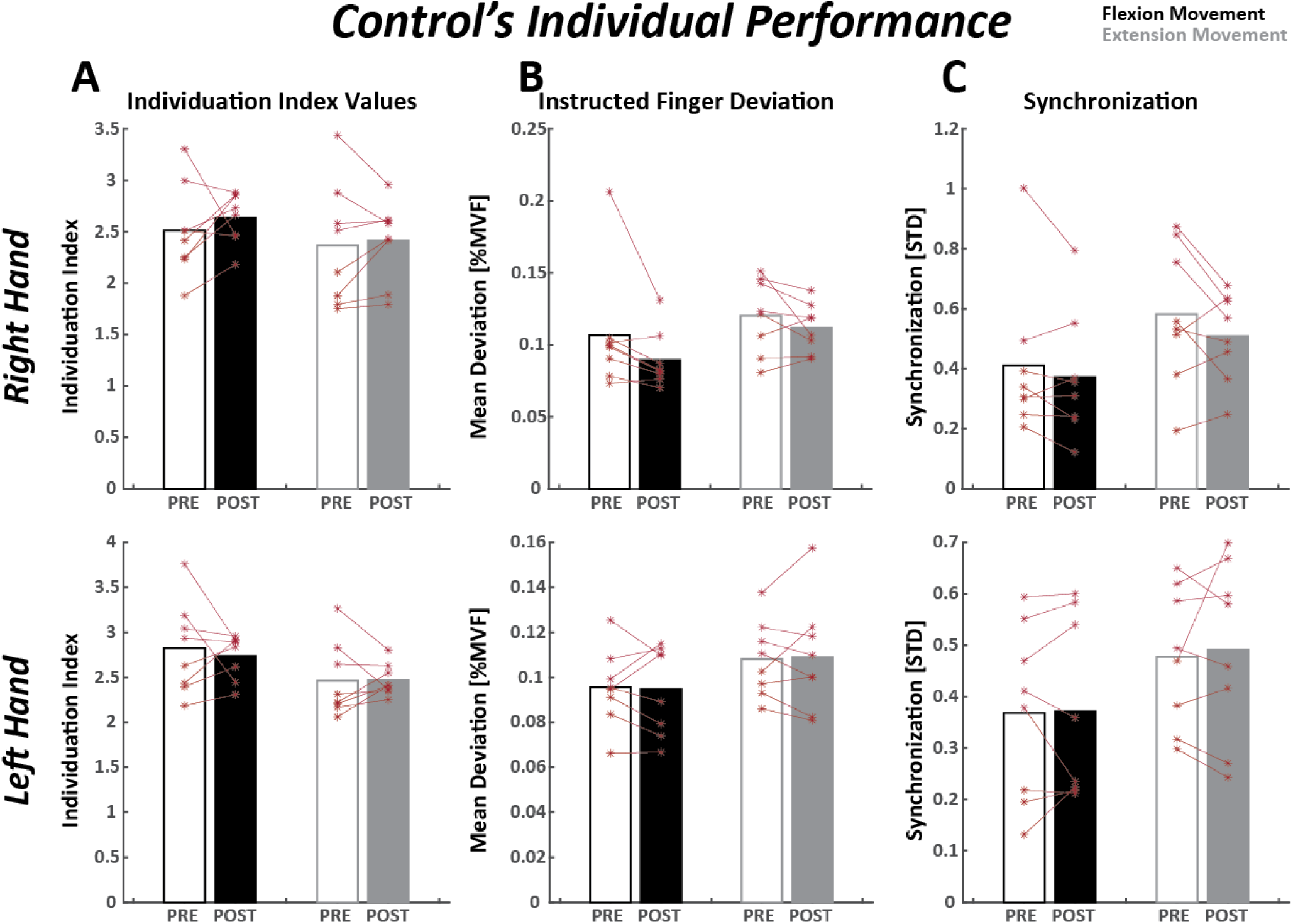
– Individual data of the control group’s performance. Each participant is represented by two connected asterisks which show the change in performance between the Pre and Post days. Flexion motion is depicted in black and extension motion is depicted in grey. There is no distinct trend among the different participants in any of the metrics. (A) The individuation index of the individual participants can be seen for both directions for right hand (top) and left hand (bottom) movements. (B) The synchronization of the individual participants can be seen for both directions for right hand (top) and left hand (bottom) movements. (C) The instructed finger deviation, normalized by the MVF, of the individual participants can be seen for both directions for right hand (top) and left hand (bottom) movements

**Supplementary Figure 6.**
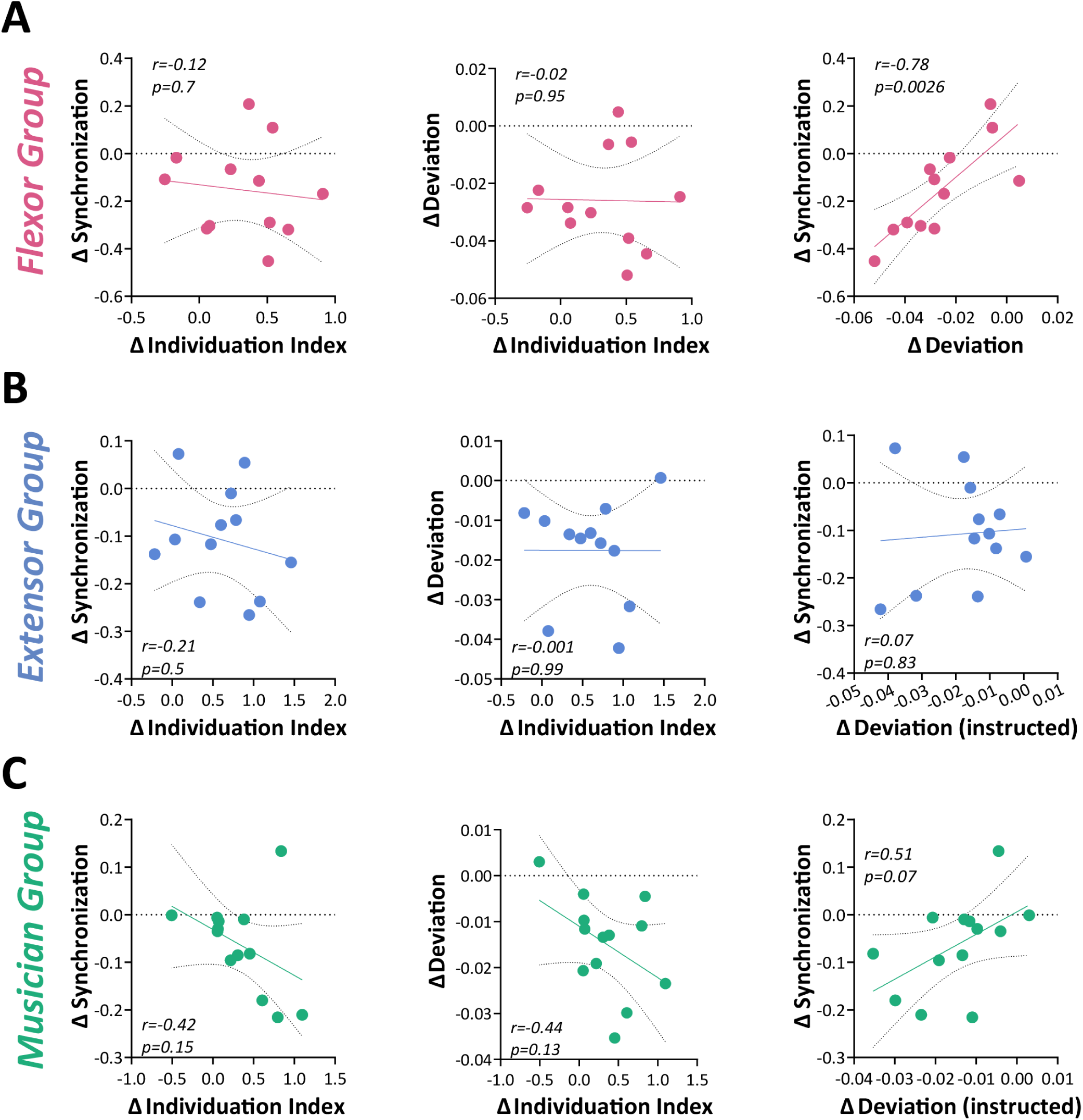
– Correlations between the synchronization, Individuation Index, and instructed finger deviation metrics for each of the three groups. The metrics are represented by the change between the values obtained pre- and post-training. Each marker represents an individual’s performance. (A) The flexor group’s correlation graphs, for right hand flexion motions. (B) The extensor groups correlation graphs, for right hand extension motions. (C) The musician groups correlation graphs, for right hand extension motions.

## Notes

### Competing Interest Statement

The authors have declared no competing interest.

## References

1. Criscimagna-Hemminger, S. E., Donchin, O., Gazzaniga, M. S. & Shadmehr, R. Learned dynamics of reaching movements generalize from dominant to nondominant arm. J. Neurophysiol. 89, 168–176 (2003).

2. Verstynen, T., Diedrichsen, J., Albert, N., Aparicio, P. & Ivry, R. B. Ipsilateral motor cortex activity during unimanual hand movements relates to task complexity. J. Neurophysiol. 93, 1209–1222 (2005).

3. Choi, J. T. & Bastian, A. J. Adaptation reveals independent control networks for human walking. Nat. Neurosci. 10, 1055–1062 (2007).

4. Huber, L. et al. Sub-millimeter fMRI reveals multiple topographical digit representations that form action maps in human motor cortex. Neuroimage 208, (2020).

5. Pruszynski, J. A. et al. Primary motor cortex underlies multi-joint integration for fast feedback control. Nature 478, 387–390 (2011).

6. Roux, F. E., Niare, M., Charni, S., Giussani, C. & Durand, J. B. Functional architecture of the motor homunculus detected by electrostimulation. J. Physiol. 598, 5487–5504 (2020).

7. Kamper, D. G., Harvey, R. L., Suresh, S. & Rymer, W. Z. Relative contributions of neural mechanisms versus muscle mechanics in promoting finger extension deficits following stroke. Muscle and Nerve 28, 309–318 (2003).

8. Divekar, N. V. & John, L. R. Neurophysiological, behavioural and perceptual differences between wrist flexion and extension related to sensorimotor monitoring as shown by corticomuscular coherence. Clin. Neurophysiol. 124, 136–147 (2013).

9. Zaaimi, B., Edgley, S. A., Soteropoulos, D. S. & Baker, S. N. Changes in descending motor pathway connectivity after corticospinal tract lesion in macaque monkey. Brain 135, 2277–2289 (2012).

10. Cauraugh, J., Light, K., Kim, S., Thigpen, M. & Behrman, A. Chronic Motor Dysfunction After Stroke. Stroke 31, 1360–1364 (2000).

11. Fritz, S. L., Light, K. E., Patterson, T. S., Behrman, A. L. & Davis, S. B. Active finger extension predicts outcomes after constraint-induced movement therapy for individuals with hemiparesis after stroke. Stroke 36, 1172–1177 (2005).

12. Twitchell, T. E. The restoration of motor function following hemiplegia in man. Brain 74, 443–480 (1951).

13. Arbuckle, S. A. et al. Structure of population activity in primary motor cortex for single finger flexion and extension. bioRxiv 2020.03.17.996124 (2020) doi:10.1101/2020.03.17.996124.

14. Aoki, T., Furuya, S. & Kinoshita, H. Finger-Tapping Ability in Male and Female Pianists and Nonmusician Controls. Motor Control 9, 23–39 (2005).

15. Slobounov, S., Johnston, J., Chiang, H. & Ray, W. The role of sub-maximal force production in the enslaving phenomenon. Brain Res. 954, 212–219 (2002).

16. Zatsiorsky, V. M., Li, Z. M. & Latash, M. L. Precision finger pressing force sensing in the pianist-piano interaction. Exp Brain Res 187–195 (2000).

17. Lang, C. E. & Schieber, M. H. Human finger independence: Limitations due to passive mechanical coupling versus active neuromuscular control. J. Neurophysiol. 92, 2802– 2810 (2004).

18. Schieber, M. H. & Santello, M. Hand function: Peripheral and central constraints on performance. J. Appl. Physiol. 96, 2293–2300 (2004).

19. Yu, W. S., Van Duinen, H. & Gandevia, S. C. Limits to the control of the human thumb and fingers in flexion and extension. J. Neurophysiol. 103, 278–289 (2010).

20. van Duinen, H., Yu, W. S. & Gandevia, S. C. Limited ability to extend the digits of the human hand independently with extensor digitorum. J. Physiol. 587, 4799–4810 (2009).

21. Furuya, S. & Soechting, J. F. Speed invariance of independent control of finger movements in pianists. J. Neurophysiol. 108, 2060–2068 (2012).

22. Tominaga, K., Lee, A., Altenmüller, E., Miyazaki, F. & Furuya, S. Kinematic Origins of Motor Inconsistency in Expert Pianists. PLoS One 11, e0161324 (2016).

23. Oku, T. & Furuya, S. Skilful force control in expert pianists. Exp. Brain Res. *2017* 2355 235, 1603–1615 (2017).

24. Oldfield, R. C. The Assessment and Analysis of Hnadedness: The Edinburgh Inventory. Encyclopedia of Clinical Neuropsychology 1209–1209 (2011) doi:10.1007/978-0-387-79948-3_6053.

25. Brainard, D. H. The Psychophysics Toolbox. Spat. Vis. 10, 433–436 (1997).

26. Mawase, F. et al. Pushing the Rehabilitation Boundaries: Hand Motor Impairment Can Be Reduced in Chronic Stroke. Neurorehabil. Neural Repair 34, 733–745 (2020).

27. Xu, J. et al. Separable systems for recovery of finger strength and control after stroke. J. Neurophysiol. 118, 1151–1163 (2017).

28. Schieber, M. H. Individuated finger movements of rhesus monkeys: A means of quantifying the independence of the digits. J. Neurophysiol. 65, 1381–1391 (1991).

29. Zatsiorsky, V. M., Li, Z. M. & Latash, M. L. Coordinated force production in multi-finger tasks: Finger interaction and neural network modeling. Biol. Cybern. 79, 139–150 (1998).

30. Abolins, V. & Latash, M. L. The Nature of Finger Enslaving: New Results and Their Implications. Motor Control 25, 680–703 (2021).

31. Sanei, K. & Keir, P. J. Independence and control of the fingers depend on direction and contraction mode. Hum. Mov. Sci. 32, 457–471 (2013).

32. Perez, M. A. & Rothwell, J. C. Distinct Influence of Hand Posture on Cortical Activity during Human Grasping. J. Neurosci. 35, 4882–4889 (2015).

33. Stinear, C. M. & Byblow, W. D. Role of intracortical inhibition in selective hand muscle activation. J. Neurophysiol. 89, 2014–2020 (2003).

34. Beck, S. & Hallett, M. Surround inhibition in the motor system. Exp. Brain Res. *2011* 2102 210, 165–172 (2011).

35. Sohn, Y. H. & Hallett, M. Surround inhibition in human motor system. Exp. Brain Res. 2004 1584 158, 397–404 (2004).

36. Sohn, Y. H. & Hallett, M. Disturbed surround inhibition in focal hand dystonia. Ann. Neurol. 56, 595–599 (2004).

37. Waters-Metenier, S., Husain, M., Wiestler, T. & Diedrichsen, J. Bihemispheric transcranial direct current stimulation enhances effector-independent representations of motor synergy and sequence learning. J. Neurosci. 34, 1037–1050 (2014).

38. Hazeltine, E., Aparicio, P., Weinstein, A. & Ivry, R. B. Configural Response Learning: The Acquisition of a Nonpredictive Motor Skill. J. Exp. Psychol. Hum. Percept. Perform. 33, 1451–1467 (2007).

39. Morton, S. M. & Bastian, A. J. Prism adaptation during walking generalizes to reaching and requires the cerebellum. J. Neurophysiol. 92, 2497–2509 (2004).

40. Krakauer, J. W., Mazzoni, P., Ghazizadeh, A., Ravindran, R. & Shadmehr, R. Generalization of motor learning depends on the history of prior action. PLoS Biol. 4, 1798–1808 (2006).

41. Waters, S., Wiestler, T. & Diedrichsen, J. Cooperation Not Competition: Bihemispheric tDCS and fMRI Show Role for Ipsilateral Hemisphere in Motor Learning. J. Neurosci. 37, 7500–7512 (2017).

42. Kim, S. G. et al. Functional Magnetic Resonance Imaging of Motor Cortex: Hemispheric Asymmetry and Handedness. Science *(80-. ).* **261**, 615–617 (1993).

43. Kawashima, R. et al. Regional cerebral blood flow changes of cortical motor areas and prefrontal areas in humans related to ipsilateral and contralateral hand movement. Brain Res. 623, 33–40 (1993).

44. Haaland, K. Y., Elsinger, C. L., Mayer, A. R., Durgerian, S. & Rao, S. M. Motor Sequence Complexity and Performing Hand Produce Differential Patterns of Hemispheric Lateralization. J. Cogn. Neurosci. 16, 621–636 (2004).

45. Schluter, N. D., Rushworth, M. F. S., Passingham, R. E. & Mills, K. R. Temporary interference in human lateral premotor cortex suggests dominance for the selection of movements. A study using transcranial magnetic stimulation. Brain 121, 785–799 (1998).

46. Schluter, N. D., Krams, M., Rushworth, M. F. S. & Passingham, R. E. Cerebral dominance for action in the human brain: the selection of actions. Neuropsychologia 39, 105–113 (2001).

47. Zijdewind, I. & Kernell, D. Bilateral interactions during contractions of intrinsic hand muscles. J. Neurophysiol. 85, 1907–1913 (2001).

48. Shinohara, M., Keenan, K. G. & Enoka, R. M. Contralateral activity in a homologous hand muscle during voluntary contractions is greater in old adults. J. Appl. Physiol. 94, 966–974 (2003).

49. Post, M., Bakels, R. & Zijdewind, I. Inadvertent Contralateral Activity during a Sustained Unilateral Contraction Reflects the Direction of Target Movement. J. Neurosci. 29, 6353–6357 (2009).

50. Graziano, M. S. A. & Aflalo, T. N. Mapping behavioral repertoire onto the cortex. Neuron 56, 239–251 (2007).

51. Ejaz, N., Hamada, M. & Diedrichsen, J. Hand use predicts the structure of representations in sensorimotor cortex. Nat. Neurosci. 18, 1034–1040 (2015).

52. Schieber, M. H. Modern coordinates for the motor homunculus. J. Physiol. 598, 5305– 5306 (2020).

53. Furuya, S., Nakamura, A. & Nagata, N. Transfer of piano practice in fast performance of skilled finger movements. BMC Neurosci. 14, 1–8 (2013).

54. Gentner, R. et al. Encoding of Motor Skill in the Corticomuscular System of Musicians. Curr. Biol. 20, 1869–1874 (2010).

55. Hirano, M. et al. Acquisition of skilled finger movements is accompanied by reorganization of the corticospinal system. J. Neurophysiol. 119, 573–584 (2018).

56. Kimoto, Y., Hirano, M. & Furuya, S. Adaptation of the Corticomuscular and Biomechanical Systems of Pianists. Cereb. Cortex 32, 709–724 (2022).

57. Fiser, J. & Aslin, R. N. Unsupervised statistical learning of higher-order spatial structures from visual scenes. Psychol. Sci. 12, 499–504 (2001).

58. Principles of perceptual learning and development. - PsycNET. https://psycnet.apa.org/record/1969-35014-000.

59. Keele, S. W. & Posner, M. I. PROCESSING OF VISUAL FEEDBACK IN RAPID MOVEMENTS. J. Exp. Psychol. 77, 155–158 (1968).

